# Extensive uORF translation from HIV-1 transcripts conditions DDX3 dependency for expression of main ORFs and elicits specific T cell immune responses in infected individuals

**DOI:** 10.1101/2022.04.29.489990

**Authors:** Emmanuel Labaronne, Didier Décimo, Lisa Bertrand, Laura Guiguettaz, Thibault J.M. Sohier, David Cluet, Valérie Vivet-Boubou, Clara Dahoui, Pauline François, Isabelle Hatin, Olivier Lambotte, Assia Samri, Brigitte Autran, Lucie Etienne, Caroline Goujon, Jean-Christophe Paillart, Olivier Namy, Berta Cecilia Ramirez, Théophile Ohlmann, Arnaud Moris, Emiliano P. Ricci

**Author notes:** Equal contribution (Didier Décimo, Lisa Bertrand).

## Abstract

Human immunodeficiency virus type-1 (HIV-1) is a complex retrovirus which relies on alternative splicing, translational and post-translational mechanisms to produce more than 15 functional proteins from its single ∼10kb transcriptional unit. Here, we have applied ribosome profiling and nascent protein labeling at different time points during infection of CD4+ T lymphocytes to characterize the translational landscape of cellular and viral transcripts during the course of infection. Our results indicate a strong impact of viral infection on host cellular transcript levels but a modest impact on global translation rates. Analysis of ribosome profiling reads from viral transcripts reveals extensive and productive non-AUG translation of small peptides from multiple upstream open reading-frames (uORFs) located in the 5’ long terminal repeat. Remarkably, these uORFs derived peptides elicit specific T cell responses in HIV-infected individuals. uORFs are conserved among other retroviruses and, together with the TAR sequence, condition the dependency on DDX3 for efficient translation of the main viral open-reading frames.

## Introduction

HIV-1 is a single-stranded, positive-sense, diploid and enveloped RNA virus that belongs to the Retroviridae family. Upon infection, the genomic RNA is reverse-transcribed into a double-stranded DNA molecule that is then integrated into the host cell genome as a provirus or in some instances remains unintegrated in a circular form. The provirus DNA (integrated or circular) is transcribed by host RNA polymerase II (Pol II) into a single capped and polyadenylated RNA. Like other complex retroviruses, this single RNA species undergoes extensive alternative splicing to produce a multitude of matured transcripts coding for regulatory proteins in addition to the canonical Gag, Pol and Env proteins (Emery et al., 2017; Nguyen Quang et al., 2020; Ocwieja et al., 2012). Alternative splicing of the viral transcript and expression of the different viral proteins is temporally regulated during the course of infection. Fully spliced transcripts coding for regulatory proteins are preferentially exported to the cytoplasm during early stages of the replication cycle. Partially spliced mRNAs and the unspliced genomic RNA (the latter also coding for Gag and Gag-Pol) are preferentially exported at late stages of the replication cycle through a process that requires the viral encoded protein Rev (Felber et al., 1989).

Translation of viral transcripts is also tightly regulated during the replication cycle through multiple mechanisms of translation initiation for both the unspliced genomic RNA and several spliced variants (de Breyne and Ohlmann, 2018; Guerrero et al., 2015; Jaafar and Kieft, 2019). Indeed, the 5’ untranslated region (UTR) of viral transcripts is relatively long (>300 nucleotides) and contains several extensive RNA structures (such as TAR, Poly(A), PBS and the packaging signal stem-loops) that are essential at different steps of the replication cycle but may represent an obstacle for 43S ribosomes to bind viral mRNAs and reach the translation start codon (Ricci et al., 2008a). As such, internal ribosome entry sites (IRESes) have been described in the 5’UTR of unspliced and some spliced viral transcripts as well as in the coding region of Gag for unspliced RNAs (Brasey et al., 2003; de Breyne and Ohlmann, 2018; Daudé et al., 2016; Deforges et al., 2017; Herbreteau et al., 2005; Locker et al., 2011; Plank et al., 2013; Ricci et al., 2008b; Vallejos et al., 2012). Furthermore, the viral protease encoded from the full-length RNA is capable of mediating proteolytic cleavage of the translation initiation factors eIF4G, eIF4G2 and PABP, which inhibits cap-dependent translation and 43S ribosomal scanning (Alvarez et al., 2003; Castelló et al., 2009; Ohlmann et al., 2002; Prévôt et al., 2003; Ricci et al., 2008b; Ventoso et al., 2001). Translation initiation on viral RNAs can also occur through a cap-dependent process that is facilitated by many host proteins including RNA helicases and RNA binding proteins that facilitate recruitment and scanning of 40S on viral transcripts (Bolinger et al., 2010; de Breyne and Ohlmann, 2018; Chatel-Chaix et al., 2004; Dugré-Brisson et al., 2004; García-de-Gracia et al., 2021; Hartman et al., 2006; Ramos et al., 2021; Singh et al., 2022; Soto-Rifo et al., 2012, 2013). Translation of the full-length RNA is also regulated at the elongation step, through a frameshifting signal which allows the expression of the structural Gag polyprotein and the fusion Gag-Pol polyprotein that contains viral proteins with enzymatic activity such as the protease, reverse-transcriptase and integrase (Jacks et al., 1988).

The regulation of viral protein translation has a strong influence on the initiation of immune responses to the virus, specifically on T cell mediated immunity. T cells recognize, on infected cells, virus-derived peptides presented by major histocompatibility complex (MHC) molecules. In particular, CD8+ T cells recognize viral peptide, presented by MHC class-I molecules, that are derived from the processing, by proteasomes, of the native viral proteins and/or, misfolded or truncated viral polypeptides (Wei et al., 2019; Yewdell, 2020). There is a direct link between translation, protein degradation and the loading of MHC-I molecules (Erhard et al., 2020; Pierre, 2009; Wei et al., 2019), in particular in HIV-infected cells (Casartelli et al., 2010; Schubert et al., 2000). Interestingly, CD4+ T cells can also recognize HIV peptides, presented by MHC class-II molecules, that are derived from newly synthetized viral proteins (Coulon et al., 2016).

Translation of HIV-1 and closely related lentiviruses has been mainly studied using *in vitro* translation extracts or reporter constructs that have been instrumental to uncover the many mechanisms governing HIV-1 expression. However, a global view of the expression and translation of viral transcripts during a productive replication cycle is still missing. Here, we have performed ribosome profiling and RNA sequencing from cytoplasmic extracts obtained from infected cells and cells with a latent HIV-1 infection in which the replication cycle is coordinately induced. Our results highlight the temporal regulation of expression and translation of viral and cellular transcripts and indicate a modest impact of infection on overall cellular translation rates. Ribosome profiling data reveals extensive non-AUG translation events from upstream open reading frames (uORFs) located in the 5’UTR of spliced and unspliced viral mRNAs. We further demonstrate that uORFs negatively modulate translation from the main viral ORFs. We then searched for cellular factors that might alleviate the negative effect exerted by uORFs on translation of viral proteins from canonical ORFs and identified the host DEAD-box protein DDX3, which is implicated in regulating uORF translation (Calviello et al., 2021; Chen et al., 2018a; Guenther et al., 2018; Linsalata et al., 2019) as well as translation of HIV-1 viral transcripts (Soto-Rifo et al., 2012, 2013). We demonstrate that DDX3 requirement for efficient expression of HIV-1 viral transcripts alleviates the negative effect that uORFs exert on translation of viral proteins from canonical ORFs. uORF translation from viral transcripts also occurs in the closely related lentivirus HIV-2 and in the more phylogenetically distant retrovirus HTLV-1 (Human T Lymphotropic Virus 1), suggesting that it may be a conserved feature among retroviruses. Finally, using IFNγ ELISPOT assays in PBMCs from HIV-1 infected individuals, we can detect specific T cell responses directed against uORF-derived peptides thus indicating that uORFs encode MHC-ligands with potential relevance in mediating an immune response against infected cells *in vivo*.

## Results

### HIV-1 infection induces changes in abundance and translation of specific host mRNAs

Infection with HIV-1 has been previously proposed to down-regulate cap-dependent translation through the cleavage of translation initiation factors and the arrest of infected cells in G2-M phase (Alvarez et al., 2003; Castelló et al., 2009; Ohlmann et al., 2002; Sharma et al., 2012; Ventoso et al., 2001). However, this has mainly been tested under over-expression of the viral protease or transfection of plasmids coding for the infectious provirus. A global assessment of cellular and viral translation at the transcriptome-wide scale under productive infection is still missing. To monitor the impact of HIV-1 infection on gene expression, we infected SupT1 CD4+ T cells with HIV-1 (NL4-3 strain) to obtain more than 90% of cells expressing p24, 24 hours after infection, as measured by flow cytometry analysis (Supplementary Figure 1A). Mock-infected cells were grown in parallel as a control. Cytoplasmic lysates from HIV-1 and mock-infected cells were recovered at 0, 1, 12, 24 and 36 hours post infection (hpi) to monitor changes in cytoplasmic transcript abundance and ribosome loading by RNA-seq and Ribosome Profiling (Ribo-seq), respectively (Figure 1A). For time course experiments, micrococcal nuclease was used to obtain ribosome footprints instead of RNase I, which has been shown to lead to ribosome degradation (Cenik et al., 2015; Gerashchenko and Gladyshev, 2017; Ricci et al., 2014). However, because tri-nucleotide periodicity is not as clear when working with micrococcal nuclease, samples at 24hpi were also prepared with RNase I in order to track the reading frame of ribosomes more accurately (Supplementary figure 1B).

**Figure 1.**
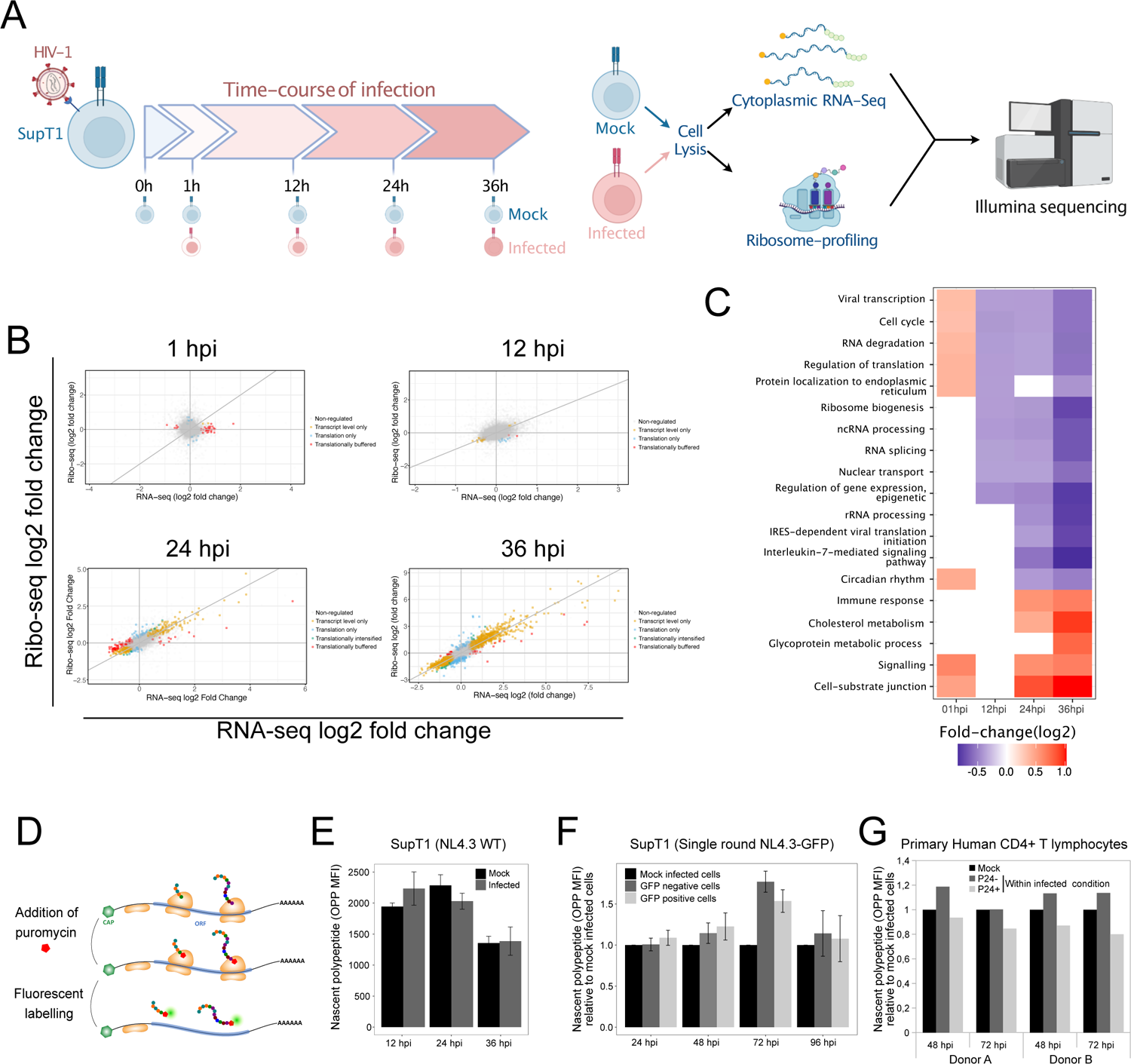
Transcriptional and translational changes in HIV-1 infected cells. **A.** Schematic representation of the procedure to monitor transcript abundance and translation in HIV-1 infected cells. Briefly, SupT1 cells were infected or not (Mock) with HIV-1 (NL4.3 strain) at MOI 5. At 0, 1, 12, 24 and 36 hours post infection (hpi), cells were lysed to recover the cytoplasmic fraction and prepare ribosome profiling and RNA-seq libraries subjected to high-throughput sequencing on the Illumina Hiseq platform. **B.** Scatter-plot of the fold-change (log2) in cytoplasmic RNA-seq and Ribo-Seq of the Mock-infected and HIV-1 infected cells at each time point of infection. Orange dots (“Transcript-level only”) corresponds to genes exclusively regulated at the transcript abundance level. Blue dots (“Translation only”) correspond to genes which display differences in ribosome occupancy while transcript abundance remains unchanged. Green dots (“Translationally intensified”) correspond to genes with significant changes in transcript abundance and significantly further changes, in the same direction, in ribosome occupancy upon infection. Red dots (“Translationally buffered”) correspond to transcripts displaying significant changes in transcript abundance but for which there is compensation at the translational level to maintain unchanged ribosome occupancy levels upon infection. **C.** Gene ontology analysis of differentially expressed genes at each time point. **D.** (Left panel) Scheme describing nascent protein labeling using O-propargyl-puromycin (OPP). Briefly, cells are incubated with OPP and fixed in paraformaldehyde before a fluorophore is covalently linked through a click-reaction. Cells are then analyzed by flow cytometry to monitor signal intensity at a single-cell level. **E.** Flow cytometry analysis of OPP signal in SupT1 cells infected with HIV-1, 12, 24 and 36 hpi. **F.** OPP signal in SupT1 cells infected with single-round recombinant HIV-1 virions coding for GFP at 24, 48, 72 and 96 hpi. **G.** OPP signal in primary Human CD4+ T cells infected with HIV-1, 48 and 72 hpi and either positive or negative for p24 expression (p24+ or p24-). Results from two independent donors are displayed separately for each time point tested.

Comparison of RNA-seq and Ribo-seq reads at 1hpi indicates that infection induces a mild (less than two-fold) but significant change in the abundance of a handful of cellular transcripts (Figure 1B, Top-left panel). However, their ribosome-loading is not affected, indicating that they have probably not engaged efficiently with the translation machinery or that the cell is buffering their expression at the translational level. At 12hpi, the overall distribution of RNA-seq and Ribo-seq reads correlates well and indicates global but mild (mostly not-significant) changes in transcript abundance (Figure 1B, Top-right panel). At 24 and 36hpi, we observed a strong and global effect on gene expression, with thousands of cellular transcripts being differentially expressed (Figure 1B, Bottom panels). Surprisingly, most changes are driven at the level of cytoplasmic transcript abundance (which can result from changes in transcription, nuclear export or transcript stability), with only a minor subset of genes (423 in total) being regulated at the translational level 36hpi.

To perform differential gene expression and gene-ontology analysis, we further used ribosome profiling reads. Indeed, these correspond to the combination of transcript abundance and ribosome loading and should therefore better correlate with protein output than RNA-seq reads. Supporting this hypothesis, we found that quantifying gene expression with Ribosome profiling data correlates better with proteomics data of HIV-infected SupT1 cells (Golumbeanu et al., 2019) than RNAseq datasets (Supplementary Figure 1C). Gene ontology analysis on up-regulated genes at 36hpi shows an enrichment for proteins involved in cell communication, signaling, immune system processes, and cholesterol pathway, among others (Figure 1C). Down-regulated genes are enriched in RNA splicing, rRNA processing, ribosomal large and small subunit biogenesis, interleukin-7-mediated pathway, cell cycle, regulation of translation, IRES-dependent viral translation initiation (Figure 1C). Gene ontology analysis performed on differentially translated transcripts (423 genes in total 36hpi, 301 up-regulated and 122 down-regulated) did not show any enrichment of specific functional category. Nevertheless, down-translated genes have significantly lower GC content in their CDS and longer CDS than control genes and vice versa for up-translated genes (Supplementary Figure 1D). This suggests a possible role of the endogenous host restriction factor Schlafen 11 (SLFN11), which inhibits HIV-1 replication through GC content and codon-usage (Li et al., 2012). Further analysis of the abundance of differentially expressed transcripts across all time-points indicate that most genes display a gradual trajectory, with mild but coherent changes in cytoplasmic abundance at 12hpi, which become more pronounced as infection progresses (Supplementary Figure 1E).

### Global translation rates are modestly affected by HIV-1 infection

In the absence of synthetic exogenous RNA spike-in controls, high-throughput sequencing approaches, such as RNA-seq and ribosome profiling, can inform about relative changes in gene expression but are not able to accurately monitor bulk changes in intracellular transcript abundance or ribosome occupancy (Jiang et al., 2011). To test whether HIV-1 infection is accompanied by changes in overall translation efficiency in infected cells, we probed absolute levels of protein synthesis in infected cells using O-propargyl-puromycin (OPP), a clickable puromycin analog that incorporates into elongating polypeptide chains (Liu et al., 2012) (Figure 1D). For this, SupT1 cells were mock-infected or infected with HIV-1 and OPP incorporation measured at 12, 24 and 36hpi (Figure 1E). As a positive control, we infected HEK293T cells with Sindbis virus (SINV), an alphavirus that has been reported to induce a strong translation shutoff in host infected cells (Garcia-Moreno et al., 2019). As expected, SINV infection induced a strong decrease in OPP incorporation as soon as 12hpi (Supplementary Figure 2A, right panel). However, no significant differences in OPP incorporation were detected between mock-infected and HIV-infected cells at any of the tested time points (Figure 1E). A slight decrease of translation was nevertheless detected for both mock-infected and HIV-1 infected cells at 36hpi, probably due to the higher density of cells in the culture well.

To further validate these results at later time points of infection, SupT1 cells were infected at MOI 0.5 with VSVg pseudotyped, genetically modified single-round HIV-1 virions in which there is a frameshift mutation in the envelope coding sequence and the GFP coding sequence is placed upstream of Nef. This strategy allows to track infected cells and avoid the important cell mortality that is associated with the replication and spreading of wild-type HIV-1 virions at longer time-points of infection. To monitor translation and infection, the level of OPP incorporation and GFP expression was measured at 24, 36, 48, 72 and 96hpi. As observed (Figure 1F), translation rates in GFP positive cells (indicative of HIV-1 replication) increased slightly at 24h compared to the mock-infected control and to the GFP negative cells from the infected condition (corresponding either to non-infected cells or cells in which expression of viral proteins was still not detected). At 48 and 72h after infection, translation efficiency increased both for GFP positive and negative cells from the infected condition compared to mock-infected cells, while at 96h post-infection translation efficiency was similar in all tested conditions.

Finally, we infected activated primary CD4+ T-cells obtained from two different donors with wild-type HIV-1 virions (NL4.3 strain) and monitored translation rates at 48 and 72hpi among p24-positive and -negative cells (Figure 1G). In this set-up, we were able to detect a modest decrease in OPP incorporation (close to 20%) in the former compared to the latter, at both time points and for both donors.

Taken together, while no global translational shut-off was detected upon infection, we observed important changes in transcript abundance of host mRNAs and a milder effect on translation of specific host mRNAs in a GC-content dependent manner.

### Dynamics of HIV-1 cytoplasmic transcript expression

HIV-1 proviral DNA is transcribed by RNA polymerase II as a single-transcription unit that can be spliced to produce close to 100 different transcripts coding for viral proteins (Nguyen Quang et al., 2020; Ocwieja et al., 2012). These transcripts are expressed in a temporally coordinated manner to produce all accessory and core viral proteins required to efficiently assemble new viral particles. To quantify cytoplasmic levels of HIV-1 transcripts, we collected data from long read sequencing (Nanopore) of NL4-3 infected primary CD4 cells to identify all HIV-1 transcripts expressed in infected cells (Nguyen Quang et al., 2020) and used this information as a template to reconstruct isoform expression from our RNA sequencing reads. A simulated *in silico* dataset of RNA sequencing reads generated from the list of known HIV-1 transcripts, with a similar length distribution to our RNA sequencing libraries, validated our deconvolution approach (Supplementary Figure 3A). At 1hpi, we could only detect unspliced viral RNAs corresponding to incoming genomic RNAs (Figure 2A and Supplementary Figure 3B). At 12hpi, upon reverse-transcription and integration of most of the proviral DNA, viral transcripts were dominated by the fully spliced transcript coding for Nef, although we could still detect important levels of unspliced RNAs in the cytoplasm of infected cells (Figure 2A). The relative amount of fully spliced transcripts (coding for Nef, Tat, Rev) strongly decreased at 24 and 36hpi (Figure 2A). On the contrary, relative abundance of single-spliced and unspliced transcripts, which require Rev for their nuclear export (Cullen, 2003; Harris and Hope, 2000; Yedavalli et al., 2004), increased at 24 and 36hpi (Figure 2A). These results are in agreement with the known expression kinetics of viral transcripts during the replication cycle of HIV-1 (Davis et al., 1997; Kim et al., 1989).

**Figure 2.**
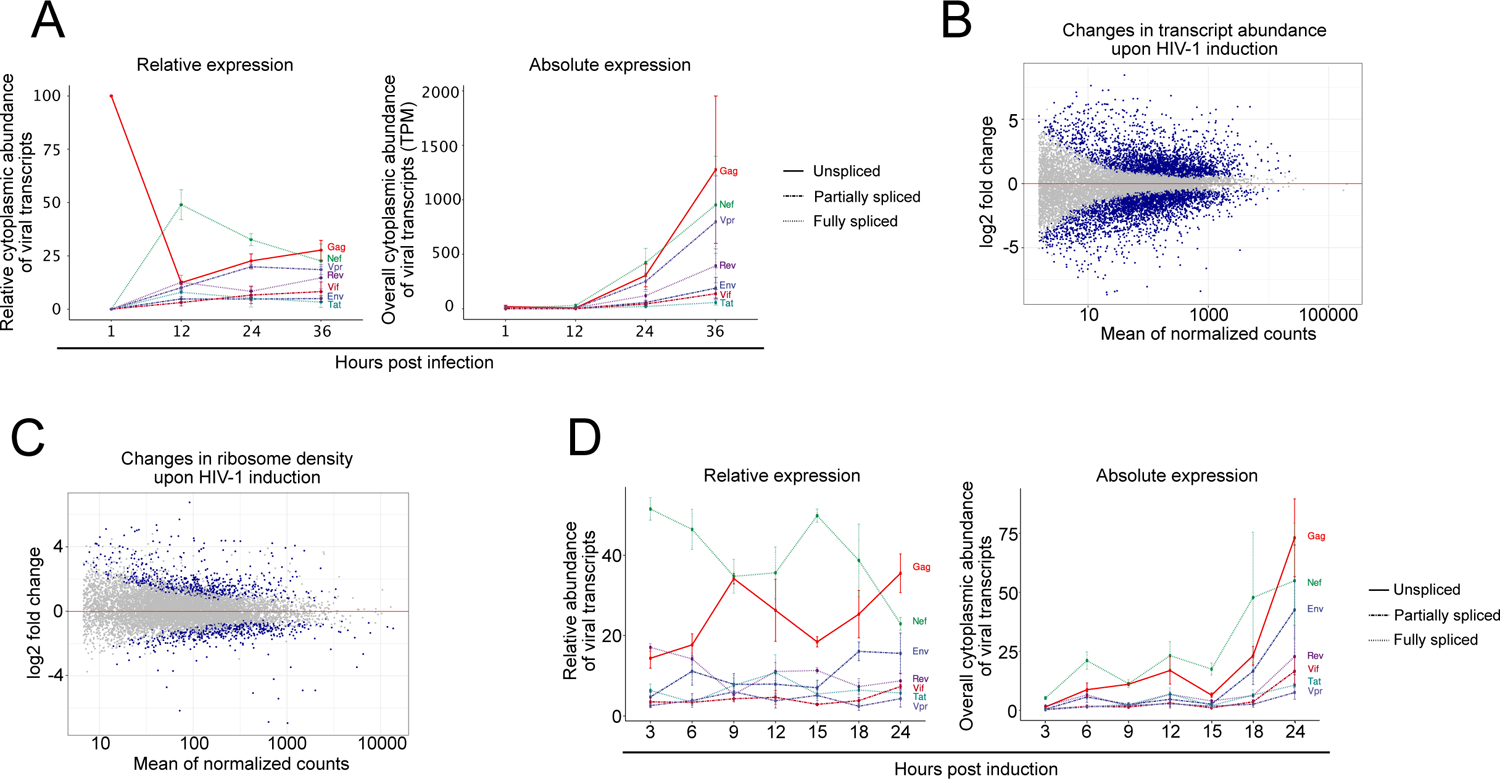
Relative and absolute expression of viral transcripts during infection. **A.** (Left panel) Relative cytoplasmic amount of viral transcripts (the sum of all transcripts at any given time point corresponding to 100%) in SupT1 cells, 1, 12, 24 and 36 hpi. (Right panel) Overall abundance of viral transcripts within RNA-seq libraries displayed in transcripts per million (TPM). **B.** MA-plot (log2 fold change Vs mean average) of cytoplasmic RNA-seq reads from U937 cells with a latent HIV-1 integrated proviral DNA, 24 hours after induction of proviral DNA expression. Blue dots correspond to differentially expressed genes with an adjusted p-value < 0.05. **C.** Same as **B.** but using ribosome density values. **D.** Relative (left panel) and overall (right panel) abundance of viral transcripts in U937 cells, 3, 6, 9, 12, 15, 18 and 24 hours after induction of proviral DNA expression.

To bypass the heterogeneity introduced by variable timing in reverse-transcription of incoming genomic RNAs, integration and transcription of proviral DNAs, we performed a time-course analysis of cytoplasmic RNA expression and ribosome profiling from U937 cells in which a stably integrated HIV-1 provirus lies in a latent form (Folks et al., 1987). In this system, expression of viral transcripts is triggered in a synchronous manner through incubation of cells with phorbol 12-myristate 13-acetate (PMA) and ionomycin. U937 with a latent integrated HIV-1 provirus were induced and cytoplasmic extracts recovered at 0, 3, 6, 9, 12, 15, 18 and 24 h to monitor RNA levels and mRNA translation. Differential gene expression analysis indicated a strong impact of PMA/ionomycin treatment on cellular gene expression and translation (Figure 2B and 2C). The overlap between differentially expressed transcripts obtained from PMA/ionomycin activated U937 cells and SupT1 cells infected with HIV-1 virions is nevertheless moderate (Supplementary figure 3C), suggesting that changes observed in U937 are mainly driven by PMA/ionomycin and not by expression of viral transcripts. Similarly to results obtained in SupT1 cells, viral transcripts were dominated by the fully spliced transcript coding for Nef at early time points (Figure 2D). Expression of unspliced genomic RNA was also abundant at early time-points and constantly increased throughout time in parallel with the decrease in Nef expression. Surprisingly, relative expression of all other viral transcripts was much lower than that of Nef and the genomic RNA and appeared comparatively constant throughout time.

**Figure 3.**
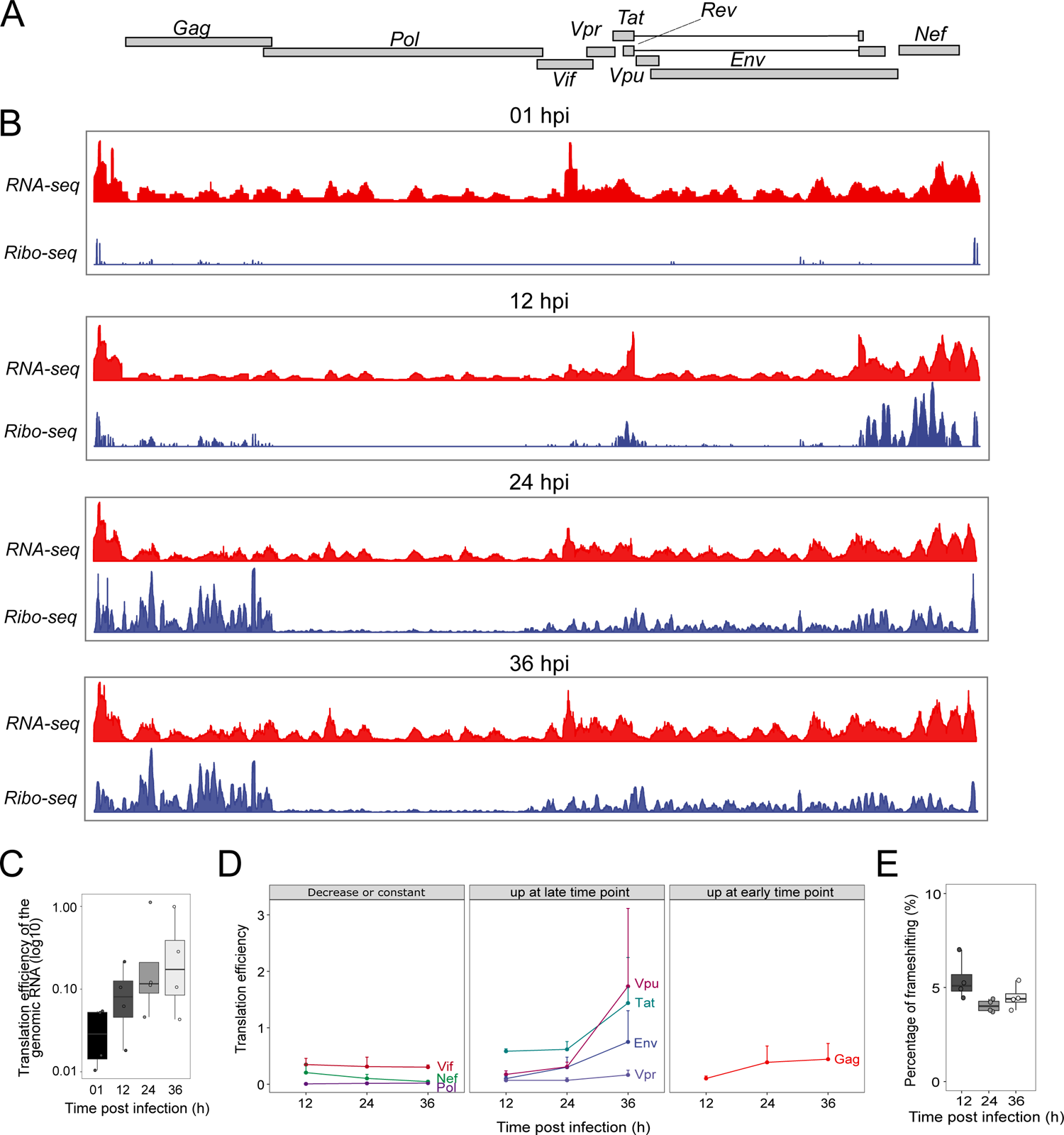
Translational landscape of viral transcripts. **A.** HIV-1 genomic structure. **B.** Distribution of RNA-seq (red) and ribosome profiling (blue) reads across the HIV-1 genome in SupT1 cells at 1, 12, 24 and 36 hpi. **C.** Translation efficiency of the genomic RNA (across both Gag and Pol coding sequences) at each time point of infection. **D.** Translation efficiency of canonical viral mRNAs, 12, 24 and 36 hpi. **E.** Percentage of Gag-Pol ribosome frameshifting at each time point of infection.

These results indicate that cytoplasmic levels of fully and partially spliced viral transcripts, as well as that of the unspliced genomic RNA are regulated in a temporal manner, as extensively described in the literature (Davis et al., 1997; Kim et al., 1989). However, expression and translation of unspliced genomic RNAs is already abundant at early time points of infection.

### Translational landscape of HIV-1 transcripts

Having characterized the dynamics of the cytoplasmic viral transcriptome, we set out to monitor its translational landscape. As described above, we could detect RNA-seq reads corresponding to cytoplasmic incoming unspliced HIV-1 genomic RNAs as soon as 1hpi (Figure 2A). Surprisingly, we could also detect ribosome footprints on viral transcripts at 1h post infection, suggesting that a fraction of incoming viral RNAs undergo translation (Figure 3A and 3B). As expected, ribosome footprints on incoming genomic RNAs were mainly located on the Gag coding sequence and very few on Pol, which depends on ribosome frameshifting for translation (Figure 3B). However, we could also detect a significant amount of ribosome profiling signal downstream of Pol, which was unexpected because unspliced mRNAs should only code for Gag and Gag-Pol. Ribosome profiling reads downstream of the Gag-Pol coding sequence could correspond to bona-fide 80S ribosomes footprints originating from alternative translation initiation mechanisms or to footprints of RNA-binding proteins co-sedimenting with 80S ribosomes. To quantify the extent of incoming genomic RNAs undergoing translation, we calculated ribosome density on the Gag and Gag-Pol coding sequences at 1, 12, 24 and 36hpi (Figure 3C). As expected, ribosome density (calculated by normalizing ribosome footprinting reads in the Gag-Pol coding region by RNA-seq reads from the same region) was significantly lower at 1h post infection than at all other time-points (2.46, 4.42 and 5.75 times lower at 1hpi than at 12, 24 and 36 hpi, respectively). This suggests that only a small fraction of incoming genomic RNAs are capable of uncoating from viral cores and undergo translation instead of reverse-transcription (Figure 3C).

At later time points, translation of viral transcripts was detected on all canonical coding sequences (Figure 3B and 3D). To quantify translation efficiency, we calculated ribosome occupancy from the non-overlapping part of each canonical open reading frame, with exception of Rev which overlaps with Tat and Nef. As observed, ribosome occupancy on Pol is much lower than in all other viral ORFs (Figure 3D), owing to its specific translation mechanism which depends on the frameshifting of ribosomes translating Gag (Jacks et al., 1988; Namy et al., 2006). Using the ratio of ribosome footprints in the Pol and Gag coding sequences, we obtained the percentage of frameshifting, which is close to 5%, in agreement with previous observations (Mouzakis et al., 2013). The percentage of frameshifting is robust in the 4∼5% range at all time points, being slightly higher at 12hpi than 24 and 36hpi (Figure 3E), suggesting that it is not subjected to strong dynamic regulation during the replication cycle.

Ribosome occupancy on viral coding sequences is however dynamically regulated during the course of infection. Ribosome occupancy on Gag increases from 12 to 24hpi and remains relatively steady at 36hpi. Ribosome occupancy on Env, Vpr and Vpu increases at 36hpi, while on the contrary that of Nef decreases from 12 to 36hpi (Figure 3D). Taken together, similarly to transcription and splicing of viral transcripts, translation of viral ORFs is tightly regulated throughout the replication cycle.

### Mapping translation initiation sites on viral transcripts

To characterize ORFs translated from HIV-1 genome and detect new ones, we use Ribocode software (Xiao et al., 2018) that allows the detection of translated ORF analyzing the periodicity of ribosome profiling signals. We performed this analysis on samples prepared using RNaseI or RNaseA+T1, and concatenate results in table 2.

To go further, we mapped translation initiation sites on viral transcripts. For that, U937 cells in which productive replication was induced by addition of PMA/ionomycin (to obtain synchronised expression of viral transcripts) for 0, 3, 6, 9, 12, 15, 18 and 24h were incubated for 15 minutes with the translation initiation inhibitor harringtonine to allow for elongating ribosomes to run-off from translating mRNAs while blocking initiating ribosomes at their translation start sites (Figure 4 and Supplementary Figure 4A) (Ingolia et al., 2011). As shown in figure 4A, this strategy led to a clear enrichment of ribosomes at the predicted start codon on host transcripts. Enrichment of ribosome profiling fragments (RPFs) at canonical translation initiation sites was also detected in viral transcripts (Figure 4B and supplementary figure 4A). The HIV-1 genomic RNA has been shown to contain an IRES that is located within the Gag coding region (downstream of the canonical Gag start codon) which allows translation of an N-terminal truncated form of Gag (also known as p40) from an in-frame AUG-start codon (Daudé et al., 2016; Deforges et al., 2017; Locker et al., 2011; Ricci et al., 2008a; Weill et al., 2010). We were able to detect a low albeit specific signal corresponding to initiating RPFs at the p40 AUG codon (Figure 4C), suggesting that the IRES is functional *in cellula* although it is orders of magnitude less efficient than cap-dependent translation within the context of infection.

**Figure 4.**
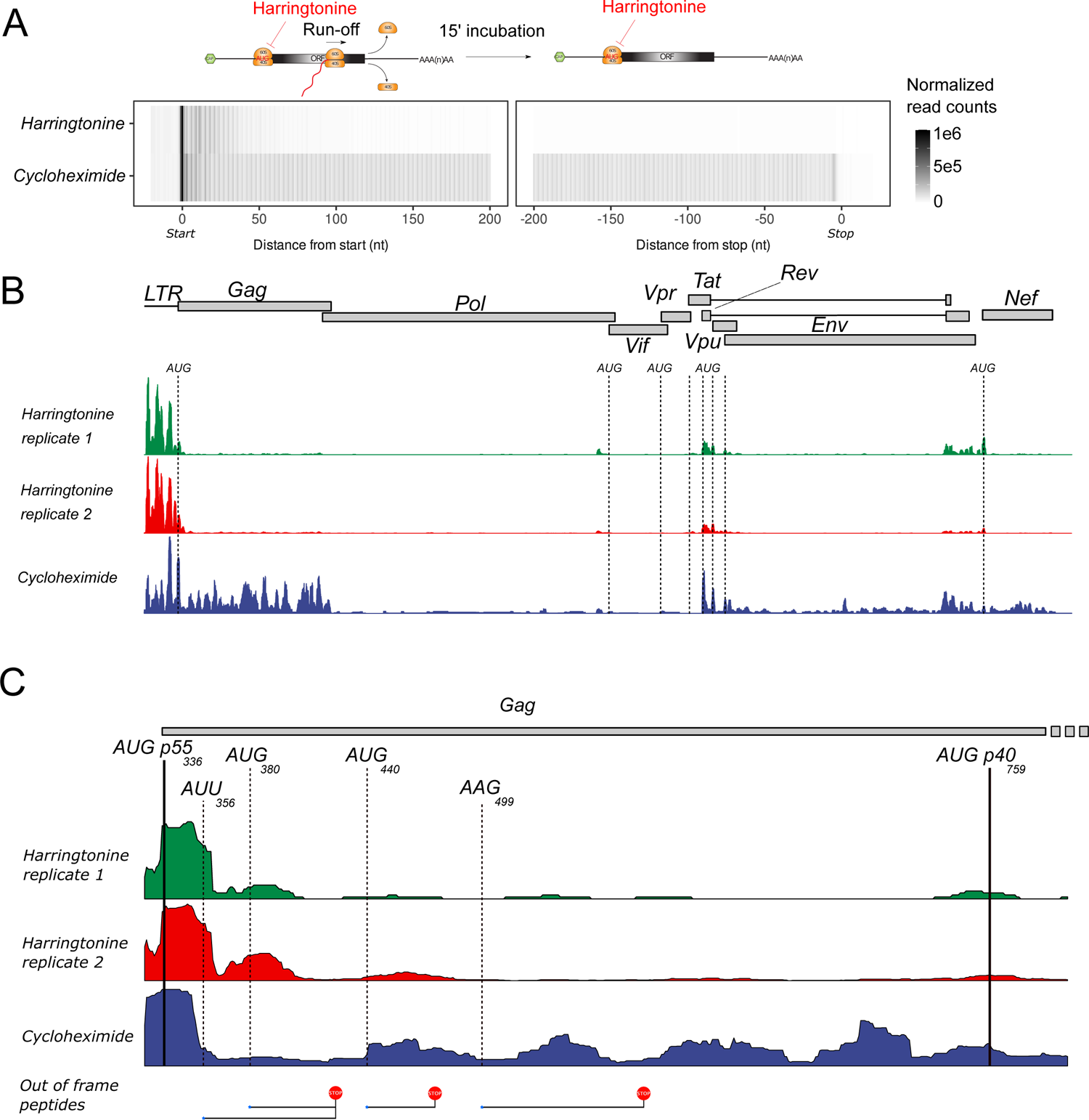
Translation initiation sites in viral transcripts. **A.** Distribution of ribosome P-sites around annotated start and stop codons in all cellular transcripts in harringtonine and cycloheximide libraries. **B.** Distribution of ribosome profiling reads across the HIV-1 genome obtained from harringtonine and cycloheximide treated cells. **C.** Distribution of ribosome profiling reads in the first 500 nucleotides of the Gag CDS obtained from harringtonine and cycloheximide treated cells. The canonical AUG start codon of Gag (p55 isoform) at position +336, as well as the position of other out-of-frame start codons predicted by Ribocode and lastly the position of the Gag (p40 isoform) start codon at position +759 are annotated in the figure.

### Multiple translation events from non-AUG start codons in the 5’UTR of viral transcripts

Although we could detect initiating RPFs on canonical AUG initiation codons, the majority of initiating ribosomes within viral transcripts originated from the LTR (Figure 4B and Supplementary Figure 4A). This signal was also observed in regular ribosome profiling samples prepared with cycloheximide (in the absence of Harringtonine) in SupT1 and in primary human CD4+ T lymphocytes (Puray-Chavez et al., 2020) infected with HIV-1 virions, as well as in U937 cells upon induction of HIV-1 transcription (Figure 5A). Careful observation of the ribosome profiling signal indicated that it overlapped with small ORFs starting with a near-cognate AUG codon. However, knowing the highly structured nature of the LTR and the fact that multiple RNA-binding proteins (including Gag) could be tightly bound to it and co-sediment with 80S ribosomes upon RNase treatment to obtain ribosome footprints, we could not exclude that these signals may originate from footprints of RNA-binding proteins other than ribosomes. Nevertheless, Fragment Length Organization Similarity Score (FLOSS) analysis of 5’UTR viral reads (Ingolia et al., 2014) from infected primary human T CD4+ lymphocytes and U937 cells, indicated that these reads had a similar length pattern than ribosome footprints mapped to the CDS of cellular coding mRNAs (Figure 5B). This, strongly suggests that reads originating from the 5’UTR of viral transcripts are bona-fide ribosome footprints.

**Figure 5.**
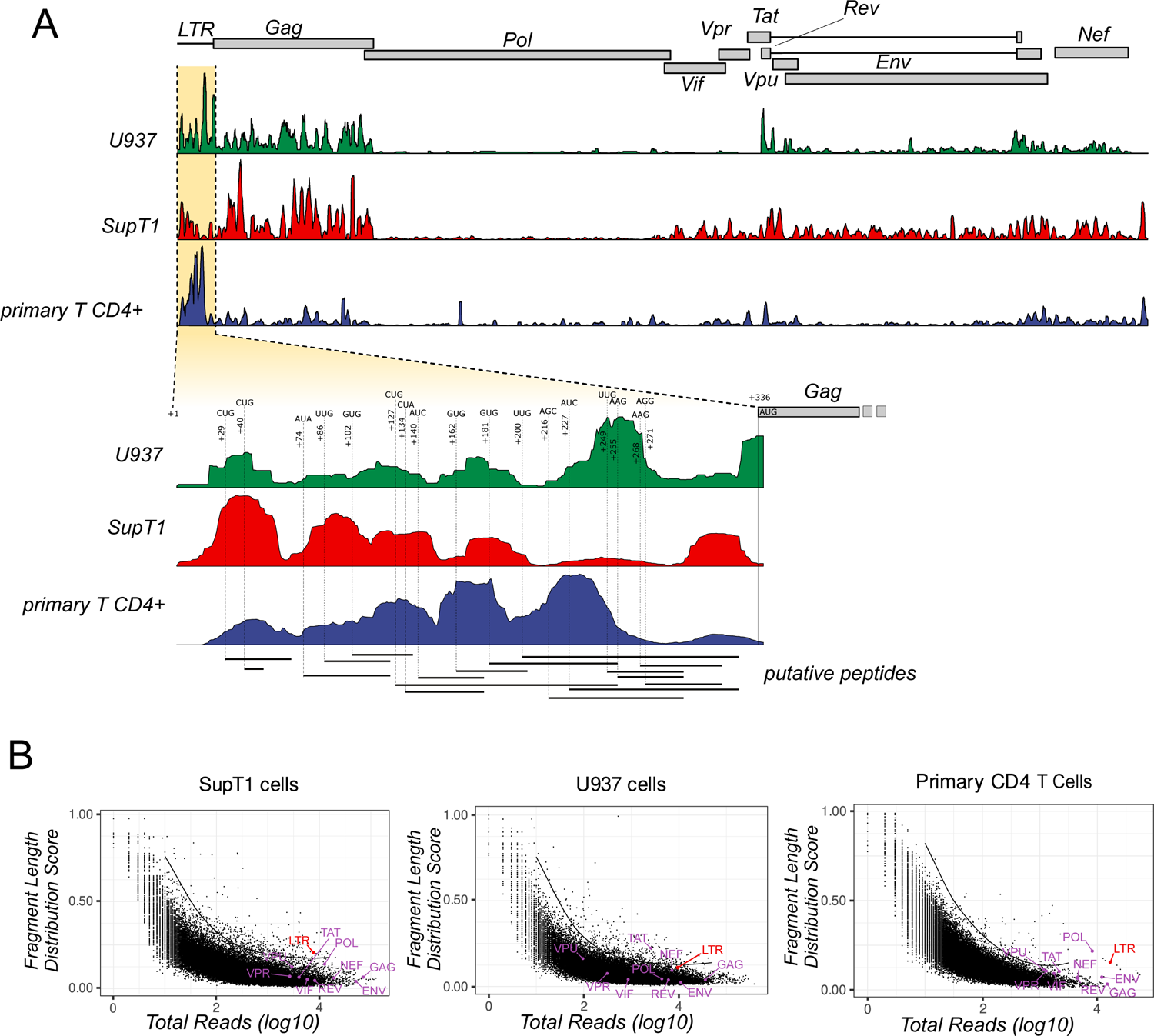
Non-AUG translation initiation sites in the 5’UTR of viral transcripts. **A.** (Top panel) Distribution of ribosome profiling reads across the HIV-1 genome from infected U937, SupT1 and primary CD4+ T lymphocytes. (Bottom panel) Close-up view of the unspliced HIV-1 5’UTR showing the distribution of ribosome profiling reads and the position of putative non-AUG start-codons. **B.** Distribution of FLOSS (Fragment Length Distribution Score) values for cellular and viral transcripts computed from ribosome profiling reads from SupT1 (left panel), U937 (middle panel) or primary CD4 T cells (right panel).

To further confirm these results, we prepared ribosome profiling libraries from HIV-1 infected cells using the ribolace kit (Clamer et al., 2018), which relies on a biotinilated puromycin analog to specifically pull-down 80S ribosomes competent for elongation. Ribosome profiling libraries were also prepared from cytoplasmic lysates incubated with 1 M of KCl prior to RNase treatment, in order to dissociate RNA-binding proteins from mRNAs as well as 80S ribosomes that are not engaged in active translation and lack a nascent peptide (Mills et al., 2016). Finally, we transfected HEK293T cells, expressing Flag-tagged ribosomes from the endogenous eL8 (RPL7a) locus, with the pNL4.3 plasmid and prepared ribosome profiling libraries by ribosome immunoprecipitation. Ribosome immunoprecipitation bypasses the use of sucrose sedimentation to recover 80S ribosomes and avoids co-sedimenting RNP complexes that could contaminate samples. As observed in Figure 6A, we could observe RPFs in the 5’UTR of viral transcripts in all tested protocols, strongly arguing for genuine uORF translation events. As a final confirmation of these results, we created reporter constructs consisting of the 5’UTR of the HIV-1 genomic RNA, a mutant version where most non-AUG initiation codons were replaced by UAG stop codons (Figure 6B, see No-uORFs 5’UTR) and a second mutant where 4 of the putative non-AUG initiation codons were replaced by AUG codons (Figure 6B, see AUG-uORFs 5’UTR), followed by the renilla luciferase coding sequence. Compensatory mutations were introduced in mutant 5’UTRs in order to maintain their overall secondary structure, which was probed by SHAPE (Gilmer et al., 2022; Merino et al., 2005; Wilkinson et al., 2006) (Supplementary Figure 5A and B). These reporter plasmids were *in vitro* transcribed to generate capped and polyadenylated RNAs that were transfected into HEK293T cells. 2h30 post-transfection, cells were collected to prepare ribosome profiling samples. As shown in Figure 6B RPFs were detected in the renilla luciferase coding sequence for all constructs tested. However, RPFs in the 5’UTR were only detected in the wild-type and AUG-uORFs mutant transcripts while almost no signal could be detected in the No-uORFs 5’UTR.

**Figure 6.**
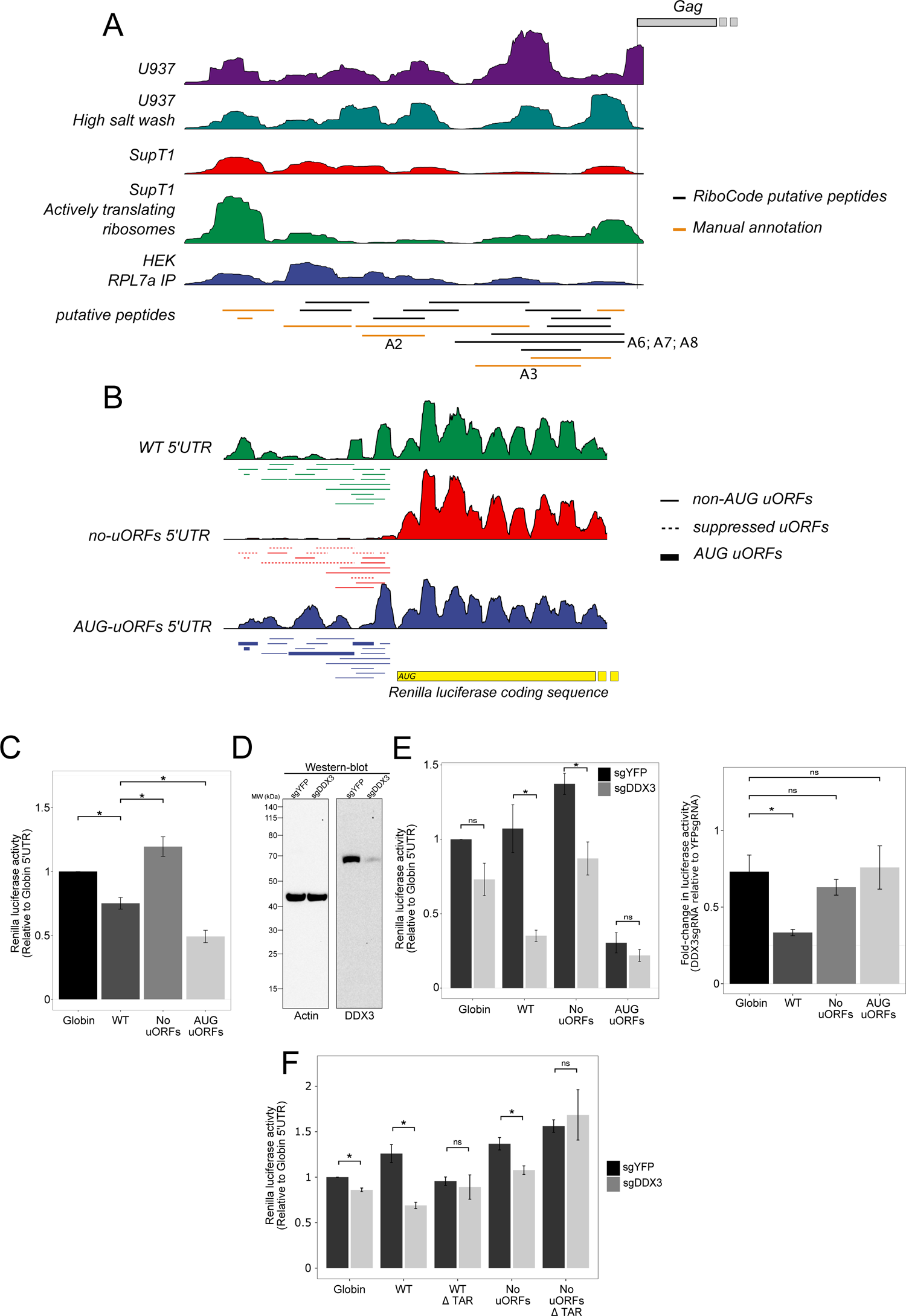
Productive translation from uORFs in viral transcripts and role of DDX3 in alleviating the negative impact of uORFs on translation from downstream main ORFs. **A.** Distribution of ribosome profiling reads across the 5’UTR of unspliced viral mRNAs in U937 cytoplasmic lysates incubated with 1M KCl (high salt wash), or from Ribolace experiments in SupT1 cells (Actively translating ribosomes), or from ribosome immunoprecipitation in transfected HEK293T cells (HEK RPL7a IP). **B.** Distribution of ribosome profiling reads on luciferase coding reporter transcripts bearing the wild-type HIV-1 5’UTR (WT 5’UTR), a mutant version in which all non-AUG uORF start codons were mutated to UAA stop codons (no uORFs 5’UTR), a mutant in which all non-AUG uORF start codons were mutated into AUG codons (AUG uORFs 5’UTR). **C.** Relative renilla luciferase activity (normalized against the Globin 5’UTR reporter mRNA) of the HIV-1 “wild-type”, “No uORFs” 5’UTR and “AUG uORFs” 5’UTR reporter mRNAs upon *in vitro* translation in the rabbit reticulocyte lysate. **D.** Western-blot analysis of β-actin (left panel) and DDX3 (right panel) expression in HEK293T cells transfected with plasmids coding for the Cas9 and a sgRNA targeting the YFP sequence (sgYFP) or the DDX3 coding sequence (sgDDX3). **E.** (Left panel) Relative renilla luciferase activity (normalized against the Globin 5’UTR reporter mRNA) from reporter mRNAs transfected in HEK293T cells in which DDX3 was knocked-down (sgDDX3) or not (sgYFP) using CRISPR-Cas9. (right panel) Fold-change of luciferase activity for each reporter mRNA between the control and DDX3 depleted condition computed from the data presented in the left panel. **F.** Relative renilla luciferase activity (normalized against the Globin 5’UTR reporter mRNA) from viral reporter mRNAs in which the TAR loop is present or absent, transfected into HEK293T cells in which DDX3 expression was knocked-down (sgDDX3) or not (sgYFP) using CRISPR-Cas9. For panels C, E and F, a Student t-test was performed to assess the differences between the mean values of compared conditions (* corresponding to a p-value <0.05).

Taken together, these results strongly suggest the presence of multiple non-AUG translation initiation sites occurring in the 5’UTR of viral transcripts that might drive productive translation of small ORF upstream of the canonical viral ORF.

### uORF translation is conserved among retroviruses

Having shown translation from uORFs originating from HIV-1 unspliced and spliced transcripts, we tested whether this was a conserved feature among retroviruses. For this, we prepared ribosome profiling from HIV-2 (Lentivirus) and HTLV-1 (Deltaretrovirus) infected cells. As shown in Supplementary figure 6C and 6D, non-AUG translation was detected at uORFs located in the 5’UTR of viral transcripts for both viruses. Interestingly, RPFs originating from non-AUG putative uORFs were also detected in the 5’UTR of Murine Leukemia Virus (Gammaretrovirus) as recently shown (Irigoyen et al., 2018) These results indicate that non-AUG translation occurring at uORFs within the 5’UTR of viral transcripts is a conserved feature of many retroviruses.

### DDX3 bypasses the negative effects of uORF on HIV-1 translation

Upstream open-reading frames generally inhibit translation of the main ORF by restricting the fraction of initiating/scanning 40S ribosomal subunits that can reach the canonical start codon (Zhang et al., 2019). To test whether uORFs located in the 5’UTR of viral transcripts restrict translation from the main ORF, we used the HIV-1 WT 5’UTR, AUG-uORFs 5’UTR and No-uORFs 5’UTR renilla luciferase reporter mRNAs described in figure 6B in addition to a control reporter mRNA bearing the 5’UTR of the human beta-globing transcript. To compare translation efficiencies from the different reporter RNAs, we first performed *in vitro* translation reactions using rabbit reticulocyte lysate (RRL). As expected, translation driven by the wild-type HIV-1 5’UTR is less efficient than that driven by the beta-globin 5’UTR in RRL (Figure 6C). Translation of the No-uORFs mutant RNA is also more efficient than that of the wild-type RNA, therefore suggesting that uORFs are able to repress translation of the main ORF by preventing scanning 40S to efficiently reach the AUG canonical codon (Figure 6C). Translation of the AUG-uORFs mutant RNA is however less efficient than that of the wild-type version, therefore suggesting that initiation on non-canonical codons is not as efficient as that occurring when they are replaced by AUG codons. The DEAD box helicase DDX3 has been shown, in yeast and humans, to mediate bypass of near-cognate start codon uORF initiation and stimulate translation from the main ORF in a large number of cellular transcripts with complex secondary structures similar to those present in the HIV-1 5’UTR (Calviello et al., 2021; Chen et al., 2018a; Guenther et al., 2018). Furthermore, efficient translation of the HIV-1 genomic RNA has been previously shown to rely on DDX3 (Soto-Rifo et al., 2012, 2013). To test whether DDX3 is involved in bypassing uORF translation from the HIV-1 5’UTR, we down-regulated DDX3 in HEK293 using CRISPR-Cas9 and transfected the reporter RNAs to test their translation. As expected, translation driven by the short beta-globin 5’UTR is not significantly sensitive to DDX3 inhibition, while that driven by the wild-type HIV-1 is strongly reduced upon DDX3 down-regulation (Figure 6D and 6E). Translation driven by the UAG-mutant was significantly more resistant to DDX3 depletion than that of the wild-type RNA thus indicating that removal of upstream non-AUG start codons from the HIV-1 5’UTR (“No-uORFs” reporter) partially removes DDX3-dependency for translation of the main ORF (Figure 6D and E). Interestingly, replacing the non-AUG start codons with AUG (“AUG-uORFs” reporter) almost completely removes the dependency on DDX3 for translation of the main ORF (Figure 6D and E). Overall these results strongly suggest that canonical HIV-1 ORF expression is regulated by viral 5’ LTR uORFs, in concert with DDX3. Finally, requirement of DDX3 for the translation of HIV-1 viral transcripts was shown to depend on the trans-activator response region (TAR) located immediately downstream the cap-structure(Soto-Rifo et al., 2012). As expected, removing TAR from our wild-type reporter RNA led to a loss of DDX3 dependency for its translation (Figure 6F). This was also the case when TAR was removed from the “No uORFs” reporter RNA therefore suggesting that TAR is required for DDX3 to modulate translation of the main ORF in a uORF-dependent manner.

### Peptides derived from 5’ LTR uORFs elicit HIV-specific T cell responses in HIV-infected individuals

To further demonstrate the existence of translation events from non-AUG start codons in the 5’UTR of HIV transcripts and test their potential role in the context of infected patients, we made used of the capacity of T cells to recognize peptide-derived from viral proteins and/or polypeptides presented by MHC molecules. HIV-specific T cells can indeed recognize peptide encoded by classical ORF but also by alternative reading frames (ARF) from HIV-1 genome (Bansal et al., 2015; Berger et al., 2010; Bet et al., 2015; Cardinaud et al., 2004, 2011). To this end, we selected 5 peptides derived from 3 different uORFs based on different parameters including ribosomal coverage, conservation among HIV isolates and more importantly on the prediction to bind a high number of HLA alleles with a high affinity. The peptides were used to screen for T cell responses in PBMCs of HIV-infected individuals. PBMCs from treated (ART) and untreated but aviremic (so called elite controllers, EC) individuals were stimulated with a pool of peptide containing the 5 uORF-derived peptides (POOL) and cultured in the presence of cytokines (IL2 and IL7) in order to expand peptide-specific memory T cells. On day 7 (not shown) and day 12 (Figure 7A-D and Supplementary Figure 6A-D), T cell responses against the POOL were assessed using IFNγ-Elispot. In addition, on day 12, except for donor EC-3, T cell responses to individual peptides of the pool (A02; A03; A06; A07; A08) were also evaluated (Figure 7A-D and Supplementary Figure 6A-D). As positive controls for T cell expansion and activation, PBMCs were also expanded with a pool of HIV Env-and Gag-derived peptides (HIV (+)) or a commercial pool of common virus-derived peptides (CEF (+)). Out of 7 donors tested, 6 reacted to HIV protein-derived peptides (HIV (+)). Note that the ART-3 donor, who did not show a significant T cell response to HIV protein-derived peptides, did however react to the CEF pool (Figure 7A-D and/or Supplementary Figure 6A-D). Remarkably, 3 individuals showed uORF-derived peptide-specific T cells responses, with the individual EC-1 reacting to peptides derived from the 3 uORF tested (Figure 7A). Depending on the peptide, the magnitude of uORF-derived peptide-specific T cells responses (IFNγ+ spots / 10^6^ PBMCs) varied and could reach plateau responses equivalent to responses induced by the pool of HIV protein-derived peptides (see peptide A08 and A02 Figure 7C and D, respectively). These results show that uORF-derived peptides elicit potent T cell responses in HIV-infected individuals. This further implies that uORFs encode polypeptides in the course of natural infection *in vivo*.

**Figure 7.**
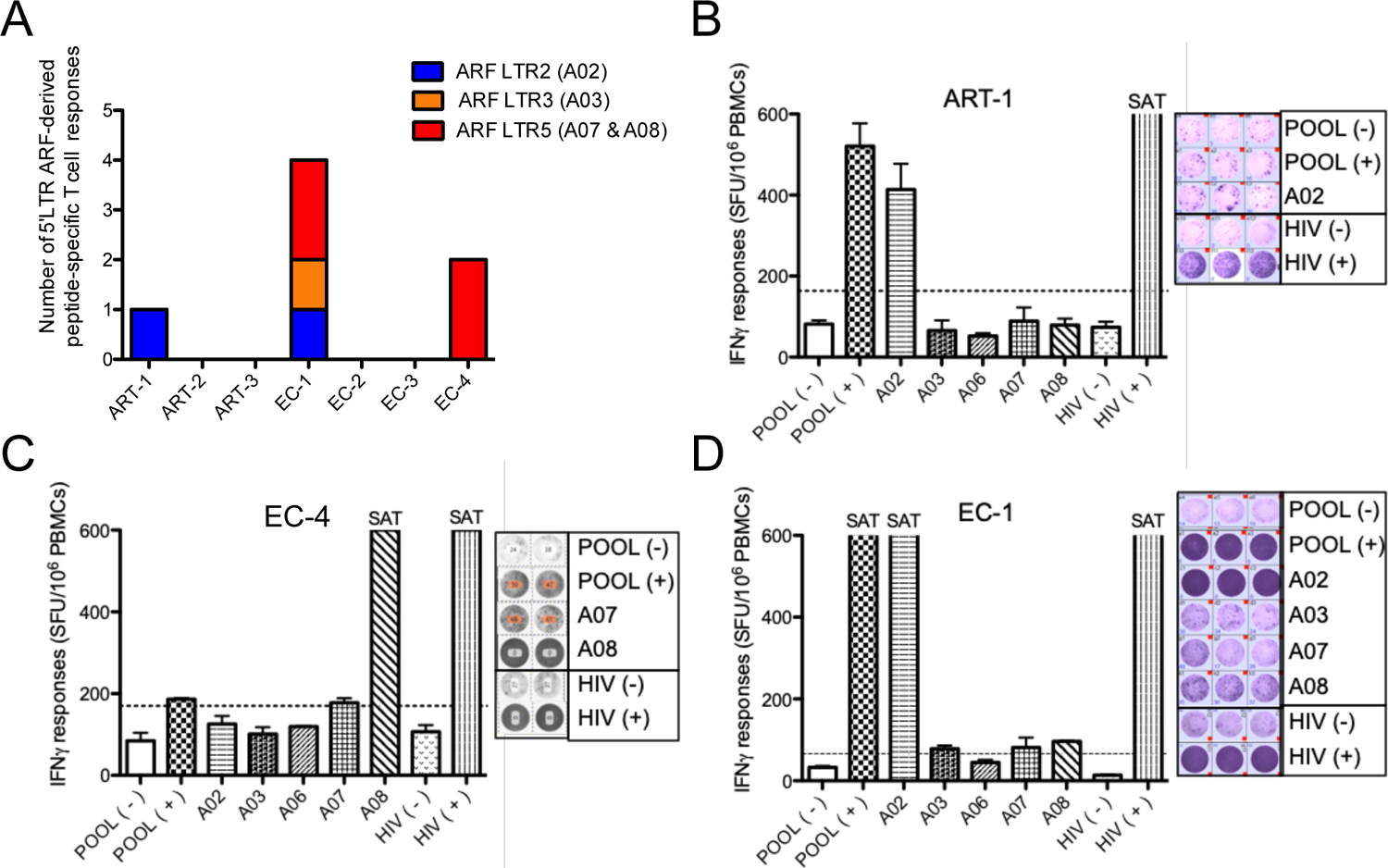
Identification of polypeptides that initiate specific T cell responses in treated and untreated HIV infected individuals. **A to D.** Based on the nucleotide sequence of NL4-3, peptides potentially encoded by uORFs (see peptides “A02; A03; A06; A07; A08” in panel **A** of this figure) of the 5’LTR were synthesized and used to screen for T cell responses in PBMCs of HIV-infected individuals. PBMCs from treated (ART) and untreated but aviremic (so called elite controllers, EC) individuals were stimulated with a pool of uORF-derived peptides (POOL) and cultured in the presence of IL-2 and IL-7 cytokines in order to expand peptide-specific memory T cells. On day 7 (not shown) and day 12, T cell responses against the POOL were assessed using IFNγ-Elispot. In addition, on day 12, except for donor EC-3, individual peptides of the pool (A02; A03; A06; A07; A08) were assessed using IFNγ-Elispot. As positive control for T cell expansion and activation, a pool of immunogenic peptides from HIV Env and Gag proteins was also used. **A.** Number of uORF-derived peptides recognized by each HIV-infected individuals. The color code indicates from which uORF the peptides are derived. **B, C and D.** left panels, detailed IFNγ-Elispot data from the 3 individuals presenting T cell responses, expressed as spot forming units (SFU) per million PBMCs; right panels pictures of the raw data from the Elispot plates in technical triplicate (**B and D**) or duplicate (**C**). POOL (-) and POOL (+): PBMCs expanded with the pool of uORF-derived peptides but restimulated on the day of the Elispot with medium or the POOL, respectively. A02; A03; A06; A07 and A08 name of individual uORF-derived peptides used for restimulation. HIV (-) and HIV (+): PBMC expanded with the pool of Env- and Gag-derived peptides but restimulated for the Elispot assay with medium or the same pool of HIV peptides, respectively. Responses were considered positive when IFNγ production was superior to 20 spots/10^6^ PBMCs and at least twofold higher than background production from cells restimulated with medium (dotted lines). SAT: saturated signal, where counts cannot be estimated due to overwhelm IFNγ secretion by activated T cells.

## Discussion

HIV-1 is a complex retrovirus capable of expressing a multitude of viral proteins from a single transcriptional unit, through a combination of alternative splicing as well as translational and post-translational mechanisms. Expression of viral proteins is tightly coordinated across time during the replication cycle (Lefebvre et al.; Nguyen Quang et al., 2020; Ocwieja et al., 2012) and it has been proposed that HIV-1 infection is associated with a translational shutoff of cellular transcripts that could favor translation of viral transcripts (Alvarez et al., 2003; Castelló et al., 2009; Ohlmann et al., 2002; Prévôt et al., 2003; Sharma et al., 2012; Ventoso et al., 2001).

Our results show that HIV-1 infection induces a strong perturbation of the cellular cytoplasmic transcriptome and a significant change in the translational status of a restricted group of transcripts, with no detectable changes in the global translation rate of the host cell. This contrasts with other RNA viruses, such as picornaviruses or alphaviruses, that induce a rapid and strong translational shut-off of cellular transcripts, which favors translation of viral transcripts. However, a lack of translational shut-off is not surprising because HIV-1 directly relies on the host transcription machinery for synthesis of viral mRNAs which are both capped and polyadenylated and can be translated in a cap-dependent manner (Berkhout et al., 2011; de Breyne and Ohlmann, 2018; Miele et al., 1996; Ricci et al., 2008a).

Cellular transcripts which expression is down-regulated upon HIV-1 infection are mainly related to the regulation of gene expression and include factors involved in ribosome biogenesis, translation, tRNA biogenesis and aminoacylation, mRNA splicing and processing, which could result from a cellular response to the stress induced by infection. However, these changes are not associated with a significant decrease in global translation rates, but rather with transcript specific changes in translation efficiency, at least within the first 36 hours of infection. In addition to the above mentioned factors, our data revealed that transcripts coding for several host proteins involved in promoting HIV-1 expression are down-regulated upon infection, these include the DEAD-Box protein DDX3 (Rao et al., 2021; Soto-Rifo et al., 2013; Yedavalli et al., 2004), FTSJ3 (Ringeard et al., 2019), RNA helicase A (Bolinger et al., 2010; Hartman et al., 2006) among others (See Table1). Cellular transcripts which expression is up-regulated upon infection are mostly associated with the plasma membrane and related to the immune response (including interferon signaling, cell migration and adhesion molecules), cellular stress response pathways and cholesterol metabolism. Interestingly, cholesterol is an important factor to recruit Gag at the plasma membrane (Dick et al., 2012; Favard et al., 2019; Nieto-Garai et al., 2021) and cholesterol depletion has been shown to reduce virion release from infected cells (Ono et al., 2007). Thus, over-expression of factors involved in cholesterol metabolism upon infection could favor Gag assembly and particle release at late time points of the replication cycle.

Cytoplasmic abundance and translation of viral transcripts as measured by RNA-seq and ribosome profiling at different times post-infection indicates a relative sequential expression of fully spliced, partially spliced and unspliced viral transcripts in infected cells (Figure 2). Nevertheless, unspliced mRNA expression and translation is still detected at early replication times. This could be partially explained by the fact that experiments were performed in a bulk population of cells with unsynchronized infection events. However, early expression of Gag/Pol could be important for the replication cycle in addition to their canonical roles at late replication steps. For instance, Integrase is known to interact with TAR RNA and Tat, and has been recently shown to regulate proviral DNA transcription at early time points of infection (Elliott et al., 2020; Liu et al., 2021; Rocchi et al., 2021; Winans and Goff, 2020).

Monitoring translation on viral transcripts revealed the presence of multiple uORFs in the 5’UTR (Figure 5A). Raw ribosome profiling signal originating from these non-AUG uORFs is as important as that from canonical viral ORFs (Figure 5A), especially in primary CD4+ T cells. This is however due to the fact that the section of the 5’UTR, which contains the uORFs is located upstream of the first splice-donor site and therefore is common to all viral transcripts. When probing translation from a single viral mRNA species (Figure 6B), ribosome profiling signal from uORFs was lower than that from the main ORF. Nonetheless, ribosome profiling signals from uORFs could be readily detected under high-salt conditions, ribosome immunoprecipitation or through the Ribolace protocol (Figure 6A). Also mutating all non-AUG start codons into UAA stop-codons led to a complete loss of ribosome profiling signal (Figure 6B), indicating that translation from uORFs might be productive. To further demonstrate that uORFs encoded polypeptides, we reveal in HIV-infected individuals the existence of T cells specific to uORF-derived peptides. Note that, using the same protocol, no uORF-derived peptide-specific T cell responses could be detected in the PBMCs of HIV-negative individuals. In addition, in HIV+ individual tested, uORF-derived peptide-specific T cells responses could reach the magnitude induced by a pool of peptide from classical HIV proteins. Overall, these results strongly argue that uORFs encode polypeptides in the course of natural infection *in vivo*.

Remarkably, removal of uORF start codons lead to a mild but significant increase in translation from the main ORF, suggesting that they exert a negative effect on viral protein expression. Nevertheless, this negative effect is partially suppressed by the DEAD-box protein DDX3 (Figure 7D). Interestingly, translation from uORFs is conserved in other retroviruses, such as HIV-2, HTLV and MLV (Supplementary figure 6C, 6D and (Irigoyen et al., 2018)). This raises a question regarding the potential roles of these uORFs. Is it a consequence of the structural and sequence constraints imposed by the multiple functional elements located in the 5’UTR (TAR, poly(A), PBS, dimerization and encapsidation signals)? Do they play a *cis-*regulatory role by restraining the number of ribosomes reaching the main ORF? Cellular mRNAs with uORFs are known to be efficiently translated under stress conditions, similar to those induced by infection (Meijer and Thomas, 2002; Morris and Geballe, 2000). Dynamic regulation of uORF translation through cellular factors could also modulate translation from canonical ORFs in a dynamic fashion during the replication cycle. HIV-1 has been described to contain IRES sequences both in its 5’UTR and in the coding region of Gag (Brasey et al., 2003; de Breyne et al., 2012; Deforges et al., 2017; Locker et al., 2011; Plank et al., 2013; Vallejos et al., 2012; Weill et al., 2010). These IRESes could bypass uORFs to maintain efficient expression of viral proteins. However, our results indicate that translation from the IRES located in the Gag coding region is inefficient in the context of infection and that addition of AUG codons in the 5’UTR strongly represses translation from the main ORF, as previously described (Berkhout et al., 2011), arguing for a limited contribution of IRES translation compared to cap-dependent translation, at least in the context of reporter transcripts.

Peptides originating from uORFs could have a biological role as described for other small peptides produced from cellular uORFs (Jayaram et al., 2021). They might also be presented by MHC class I molecules and used by the immune system to mount potent antiviral T cell responses (Antón and Yewdell, 2014; Ouspenskaia et al., 2021). We highlight here that HIV-1 uORFs can regulate the expression of canonical HIV proteins and that this can be further modulated by host factors such as DDX3. In addition, we show that uORFs induce T cell responses in treated and untreated HIV-infected individuals. In the future it will be interesting to characterize the contribution of HIV uORF-specific T cell responses in the control of viral replication and disease progression and to ask whether the initiation of uORF-specific T cell might be counteracted by cellular factors such as DDX3 or cellular stress conditions that are known to regulate uORF translation.

## Material and Methods

### Plasmids

LentiCRISPRv2 was a gift from Feng Zhang (Addgene plasmid #52961 (Sanjana et al., 2014)). pMD2.G was a gift from Didier Trono (Addgene plasmid # 12259; http://n2t.net/addgene:12259; RRID:Addgene_12259). pBlades plasmids coding for single-guide RNAs targeting the genes coding for eL8 (RPL7a), DDX3 and YFP and were constructed as described in (Mangeot et al., 2019, 2021).

### Primers and DNA sequences

Sequence of the sgRNA targeting eL8 used to introduce the Flag-tag is 5’ GTACAGCTCACTCACCATCTT 3’. Sequence of the single-stranded DNA oligo used for insertion of a Flag-tag sequence in the eL8 locus by homology directed repair is 5’ TTACCCACAATTCCCTTTCCTTTCTCTCTCCTCCCGCCGCCCAAGATGGACTACAAAGACGATGACGACAAGGTGAGTGAGCTGTAGTTCCGTGGCACTATAGCCAGGTTCCGGCTGTA T 3’.

### Cell lines and cell culture

HIV-1 latently infected U937 Cells (U1) were obtained from the AIDS reagent program (Reagent number 165) and cultured in RPMI medium supplemented with 10% fetal calf serum (FCS). Induction of productive HIV-1 replication in latently infected U937 cells was obtained by complementing the medium with 100 ng/mL of PMA and 500 ng/mL of Ionomycin. SupT1 cells were obtained from ATCC and cultured in RPMI medium supplemented with 10% fetal calf serum (FCS).

HEK293T Flag-eL8 (Rpl7a) cells cultured in DMEM medium supplemented with 10% fetal cal serum (FCS). They were produced by transfection of lentiCRISPRv1 together with pBLADES-eL8, that expresses a sgRNA targeting the eL8 locus near the translation start codon, and a single-stranded DNA donor template for homology-directed repair containing homology arms to eL8 and a Flag-tag downstream the AUG start-codon. Upon transfection, a clonal population expressing Flag-eL8 was isolated by limit dilution and screening of isolated clones.

### Primary cells

Blood from healthy donors was obtained from the Etablissement Français du Sang, under agreement n°21PLER2019-0106. Peripheral blood mononuclear cells (PBMCs) were isolated by centrifugation through a Ficoll® Paque Plus cushion (Sigma-Aldrich). Primary human T cells were purified by positive selection using CD3 MicroBeads (Miltenyi Biotec), as previously described (Goujon et al., 2013). Primary T cells were cultured in RPMI supplemented with 10% fetal bovine serum and 1% penicillin-streptomycin, and stimulated for 48h with 10 μg/ml phytohemagglutinin (PHA) (Fisher Scientific) and 50 U/mL interleukin-2 (IL-2, Miltenyi Biotec) prior to infection.

### Viral production and infection

Wild-type HIV-1 NL4-3 was produced by standard polyethylenimine transfection of HEK293T. The culture medium was changed 6h later, and virus-containing supernatants were harvested 42h later. Viral particles were filtered, purified by ultracentrifugation through a sucrose cushion (25% weight/volume in Tris-NaCl-EDTA buffer) for 75 min at 4°C and 28,000 rpm using a SW 32 TI rotor (Beckman Coulter), resuspended in serum-free DMEM medium and stored in small aliquots at −80°C. Viral particles were titrated using an HIV-1 p24Gag Alpha-Lisa kit and an Envision plate reader (Perkin Elmer). VSV-G pseudotyped HIV-1 (NL4.3) and non replicating VSV-G pseudotyped HIV-1-GFP virions were produced by co-transfecting the wild-type HIV-1 NL4-3 or HIV-1 NL4-3 Δ-Env GFP plasmids, with the pMD2.G plasmid coding for the VSV-G envelope glycoprotein.

Infection of SupT1 cells was carried out as follows. Cells were concentrated at 10 million cells per mL and incubated for 1 h at 37°C in the presence of VSV-G pseudotyped HIV-1 virions (NL4.3 strain) at MOI 5. Following this incubation, cells were diluted in RPMI medium (supplemented with 10 % FCS) to a final concentration of 1 million cells per mL and collected at the indicated time points. Cells were then analyzed with the MACSQuant VYB cytometer.

Infection of primary T cells was carried out as follows. Activated cells (see above in the “ **primary cells**” section) were plated at 250,000 cell per well in a 96-well plate and infected with 10 or 100 ng p24Gag for 4 h prior to media replacement. At 48h and 72h post-infection, supernatants were harvested for viral production quantification by Alpha-Lisa.

### Flow cytometry analysis of p24 expression in infected cells

Cells infected with HIV-1 virions (NL4.3 strain) were fixed in PBS-paraformaldehyde 4% for 20 min at room temperature and permeabilized with PBS-0.5% Triton X-100 for 20 min. Cells were then incubated with the fluorescently labeled anti-p24 antibody (Clone: KC57, Beckman Coulter, 6604667; 1/200 dilution).

### Measurement of protein synthesis using O-propargyl-puromycin (OPP)

After infection with VSVg-pseudotyped HIV-1-GFP or wild-type HIV1 virions (NL4.3 strain), cells were treated with 10 µM OPP (Immagina Biotechnology, OP001-26) for 30 min at 37°C. Then, cells were fixed in PBS-paraformaldehyde 4% for 20 min at room temperature and permeabilized with PBS-0.5% Triton X-100 for 20 min. For cells infected with HIV1 virus NL4.3 strain, a fluorescent labeling was first made with anti-p24 antibody (Clone: KC57, Beckman Coulter, 6604667; 1/200 dilution). Then, fluorescent labeling of the OPP was made with the Click-IT^TM^ Plus Alexa Fluor^TM^ 488 Picolyl Azide (Thermo Fisher Scientific, C10641), according to the manufacturer’s instructions.

For cells infected with VSVg-pseudotyped HIV-1-GFP, fluorescent labeling of the OPP was made with the Click-IT^TM^ Plus Alexa Fluor^TM^ 555 Picolyl Azide (Thermo Fisher Scientific, C10642), according to the manufacturer’s instructions. Finally, the cells were analyzed with the MACSQuant VYB cytometer.

### Ribosome profiling

Ribosome profiling samples were prepared as described in (Heyer et al., 2015) with the following modifications. At each time point of infection, 10 mL of the cell culture suspension were collected and immediately mixed with 40 mL of ice-cold 1X PBS supplemented with 120µg/mL of cycloheximide. Cells were then pelleted at 500 g for 5 min at 4°C, the supernatant was discarded and the cell pellet resuspended in 10 mL of ice-cold 1X PBS + cycloheximide (100µg/mL). Cells were pelleted again at 500 g for 5 min, the supernatant was discarded and the cell pellet lysed in 1 ml of lysis buffer (10 mM Tris-HCl, pH 7.5, 5 mM MgCl_2_, 100 mM KCl, 1% Triton X-100, 2 mM DTT, 100 μg/ml cycloheximide and 1× Protease-Inhibitor Cocktail EDTA-free (Roche)). Lysate was homogenized with a P1000 pipettor by gentle pipetting up and down for a total of eight strokes and incubated at 4°C for 10 min. The lysate was centrifuged at 1,300 g for 10 min at 4°C, the supernatant recovered and the absorbance at 260 nm measured. For ribosome footprinting, 5 A260 units of the cleared cell lysates were incubated either for 30 min at RT either with 3 µg of micrococcal nuclease (Sigma), or 300 units of RNase T1 (Fermentas) and 500 ng of RNase A (Ambion) or for 1 h at 4°C with 350 units of RNase I (Ambion). After nuclease treatment, samples were loaded on top of a 10–50% (w/v) linear sucrose gradient (20 mM HEPES-KOH, pH 7.4, 5 mM MgCl_2_, 100 mM KCl, 2 mM DTT and 100 μg/mL of cycloheximide) and centrifuged in a SW-40Ti rotor at 35,000 rpm for 2 h 40 min at 4°C. The collected 80S fraction was complemented with SDS to 1% final and proteinase K (200 μg/mL) and then incubated at 42°C for 45 min. After proteinase K treatment, RNA was extracted with one volume of phenol (pH 4.5)/chloroform/isoamyl alcohol (25:24:1). The recovered aqueous phase was supplemented with 20 μg of glycogen, 300 mM sodium acetate, pH 5.2, and 10 mM MgCl_2_. RNA was precipitated with three volumes of 100% ethanol at −20°C overnight. After a wash with 70% ethanol, RNA was resuspended in 5 μl of water and the 3′ ends dephosphorylated with PNK (New England BioLabs) in MES buffer (100 mM MES-NaOH, pH 5.5, 10 mM MgCl_2_, 10 mM β-mercaptoethanol and 300 mM NaCl) at 37°C for 3 h. Dephosphorylated RNA footprints were then resolved on a 15% acrylamide (19:1), 8 M urea denaturing gel for 1 h 30 min at 35 W and fragments ranging from 26 nt to 32 nt size-selected from the gel. Size-selected RNAs were extracted from the gel slice by overnight nutation at RT in RNA elution buffer (300 mM NaCl, and 10 mM EDTA). The recovered aqueous phase was supplemented with 20 μg of glycogen, 300 mM sodium acetate, pH 5.2, and 10 mM MgCl_2_. RNA was precipitated with three volumes of 100% ethanol at −20°C overnight. After a wash with 70% ethanol, RNA was resuspended in 5 μl of water and subjected to cDNA library construction as described below.

For ribosome profiling samples exposed to high-salt in order to induce dissociation of non-elongating 80S ribosomes, the protocol was similar to that described above with the following modification. 1 M KCl (final concentration) was added to 5 A260 units of the cleared lysates and incubated for 30 min at 4°C. After this, samples were passed through Zeba Spin desalting columns 7K MWCO (Thermo Scientific) pre-equilibrated with cell lysis buffer (10 mM Tris-HCl, pH 7.5, 5 mM MgCl2, 100 mM KCl, 1% Triton X-100, 2 mM DTT, 100 μg/mL cycloheximide and 1× Protease-Inhibitor Cocktail EDTA-free (Roche)). Then, samples were treated with nucleases as described above to obtain ribosome footprints.

Ribolace ribosome footprinting experiments were performed following the manufacturer’s protocol (Immagina BioTechnology) with the following change, RNase I was replaced with 7 µg of micrococcal nuclease (Sigma) as described above.

Ribosome immunoprecipitation-based profiling was performed as follows. Two 10cm plates of HEK293T cells expressing FLAG-eL8 (RPL7a) from the endogenous locus were transfected with the pNL4.3-HIV-1 plasmid. 24 h after transfection, medium was removed, cells were washed with ice-cold 1X PBS supplemented with cycloheximide (final concentration of 100 µg/mL) and scraped from the plates. Cells were pelleted at 500g for 5 min and lysed in 1 mL of ice-cold lysis buffer (25mM Tris pH 7.4, 150mM NaCl, 15 mM MgCl_2_, 1mM DTT, 1X cOmplete EDTA-free protease inhibitor (Roche), 100µg/mL CHX and 1% Triton X-100) for 10 min on ice. Cell debris and nuclei were removed by centrifugation at 1,300 g for 10 min at 4°C. Cytoplasmic lysate 254nm UV absorption was measured and adjusted to 25 A.U./mL and lysate was treated with 60U RNase T1 and 10 ng RNase A per A.U. for 30 min at 25°C to generate ribosome footprints. Lysate was then diluted-up to 10 mL with ice-cold lysis buffer. 150 µL of anti-FLAG agarose beads suspension (Sigma-Aldrich, A2220) were washed 3 times with 10mL lysis buffer and added to the diluted sample. Ribosome binding was allowed to proceed for 1h at 4°C on rotating wheel. Beads were then washed for a total of 5 times in 15mL tubes using 10mL of lysis buffer for 5 min at 4°C, and tubes were changed after the 1^st^ and 4t^h^ wash to minimize contaminants recovery. Beads were washed one final time in a 2 mL low-protein binding tube with 1 mL and ribosomes were eluted using 150 µL of lysis buffer supplemented with 500 µg/mL FLAG peptide (DYKDDDDK peptide - GenScript) for 1 h at 4°C. Eluates were finally digested with proteinase K (200 µg/mL final) for 45 min at 42 °C and RNAs were purified using acid Phenol:Chloroform purification. cDNA libraries were generated as described below.

### Ribosomal RNA depletion

For RNA-seq samples, cytoplasmic RNAs were depleted from ribosomal RNAs using antisense DNA oligos complementary to rRNA and RNaseH as previously described (Adiconis et al., 2013). Following DNase treatment to remove DNA oligos, rRNA-depleted RNAs were fragmented using RNA fragmentation reagent (Ambion, Cat: AM8740) for 3 min at 70°C followed by inactivation with the provided “Stop” buffer.

### cDNA library preparation for Ribosome profiling and RNA-sequencing

High-throughput sequencing libraries were prepared as described in (Heyer et al., 2015). Briefly, RNAs (either from ribosome footprints or fragmented RNAs from total cytoplasmic RNAs) were dephosphorylated at their 3’ end using PNK (New England Biolabs, Cat: M0201) in the following buffer: 100 mM Tris-HCl pH 6.5, 100 mM magnesium acetate, 5 mM β-mercaptoethanol and incubate at 37 °C during 3 h. RNA fragments with a 3′-OH were ligated to a preadenylated DNA adaptor. Then, ligated RNAs were reverse transcribed with Superscript III (Invitrogen) with a barcoded reverse-transcription primer that anneals to the preadenylated adaptor. After reverse transcription, cDNAs were resolved in a denaturing gel (10% acrylamide and 8 M urea) for 1 h and 45 min at 35 W. Gel-purified cDNAs were then circularized with CircLigase II (Lucigen, Cat: CL4115K) and PCR-amplified with Illumina’s paired-end primers 1.0 and 2.0. PCR amplicons (12-14 cycles for RNA-seq and 4-6 cycles for ribosome profiling) were gel-purified and submitted for sequencing on the Illumina HiSeq 2000 platform.

### Bioinformatic analysis

First, fastq files were demultiplexed using Flexi-splitter software. Next, adaptor sequences were trimmed from RNAseq and Riboseq reads and bad quality reads were filtered with fastp software (v0.20.1) (Chen et al., 2018b). Filtered reads were then mapped on rRNA and tRNA sequenced with bowtie2 v2.3.4.1 (Langmead and Salzberg, 2012) to remove contaminants. Non-aligned reads are then mapped against the human genome with hisat2 v2.1.0 (Kim et al., 2019). Reads that are not mapped against the human genome are mapped against HIV-1 genome with Hisat2 v2.1.0. Duplicated reads were removed using a home-made script. Deduplicated reads were used to quantify gene expression with htseq-count v0.11.2 (Anders et al., 2015). Finally, bam files were used to generate normalized bigwig files with deeptools v3.5.1 (Ramírez et al., 2016).

The periodicity of Ribo-seq experiments was checked with plastid v0.5.1 (Dunn and Weissman, 2016). To identify new ORFs from our datasets, we first mapped filtered reads against hybrid human/HIV-1 genome with STAR v2.7.3a (Dobin et al., 2013) to generate bam files with transcript coordinates. These files are then used by RiboCode v1.2.11 (Xiao et al., 2018), RiboTish v0.2.5 (Zhang et al., 2017) and RiboTricer v1.3.2 (Choudhary et al., 2020) to identify new ORFs in the HIV-1 genome. FLOSS scores were calculated with scripts from (Ingolia et al., 2014).

### Patients and Samples

EC (n=4) were recruited from the CO21 CODEX cohort implemented by the ANRS (Agence nationale de recherches sur le SIDA et les hépatites virales). PBMCs were cryopreserved at enrolment. ECs were defined as HIV-infected individuals maintaining viral loads (VL) under 400 copies of HIV RNA/mL without cART for more than 5 years. HIV-infected efficiently treated patients (ART) (n=3) were recruited at Kremlin Bicêtre Hospital. They were treated for at least 1 year (mean of 10 years) and had an undetectable viral load using standard assays. HIV-negative individuals (n=3) were anonymous blood donors (Établissement Français du Sang).

### Ethic statement

All the subjects provided their written informed consent to participate in the study. The CO21 CODEX cohort and this sub-study were funded and sponsored by ANRS and approved by the Ile de France VII Ethics Committee. The study was conducted according to the principles expressed in the Declaration of Helsinki.

### 12-day in vitro T cell amplification prior Elipsot-assay

PBMCs were thawed according to standard procedure and rested 2-3 h in IMDM (Gibco/ThermoFisher Scientific) containing 5% human AB serum (SAB, Institut Jacques Boy), recombinant human IL-2 (rhIL2, 10 U/ml, Miltenyi) and DNase (1 U/ml, New England Biolabs), washed and enumerated. 5-9 x 10^6^ PBMCs were then seeded/well of a 24-well plate in IMDM supplemented with 10 % SAB, Pen/Strep, nonessential amino acids and sodium pyruvate (all from Gibco/ThermoFisher Scientific), Nevirapine (NVP, 1,2 µM, AIDS reagent program) to inhibit potential viral replication, and poly I:C (2 µg/ml, InvivoGen) to facilitate the presentation of long peptides by antigen presenting cells (Schuhmacher et al., 2020). PBMCs were immediately loaded with peptides (Vivitide) all at 10 µg/ml except for the CEF pool (CEF extended peptide pool for human CD8 T cells, Mabtech) used at 5 µg/ml and cultured overnight at 37°C, 5% CO_2_. HIV and/or CEF peptide pools were used as positive control for the expansion of HIV Env- and Gag-specific and common anti-virus-specific T cells, respectively. On day 1 and 3, T cell media were then complemented with rhIL2 (10 U/ml) and rhIL-7 (20 ng/ml, Miltenyi), respectively, and maintained though out the culture. On days 3, 5, 7, and 9, when the cell layer was > 70% confluent, cells were splitted 1:2 before the addition of rhIL-2, rhIL-7 and NVP. On day 7, cells were harvested, counted and a fraction submitted to IFN-γ Elispot assay using the uORF-derived peptide pool and positive controls: HIV and/or CEF peptide pools. The remaining cells were maintained in culture and submitted, on day 12, to IFN-γ Elispot assay using individual uORF peptides and the controls.

### Peptide for T cell experiments

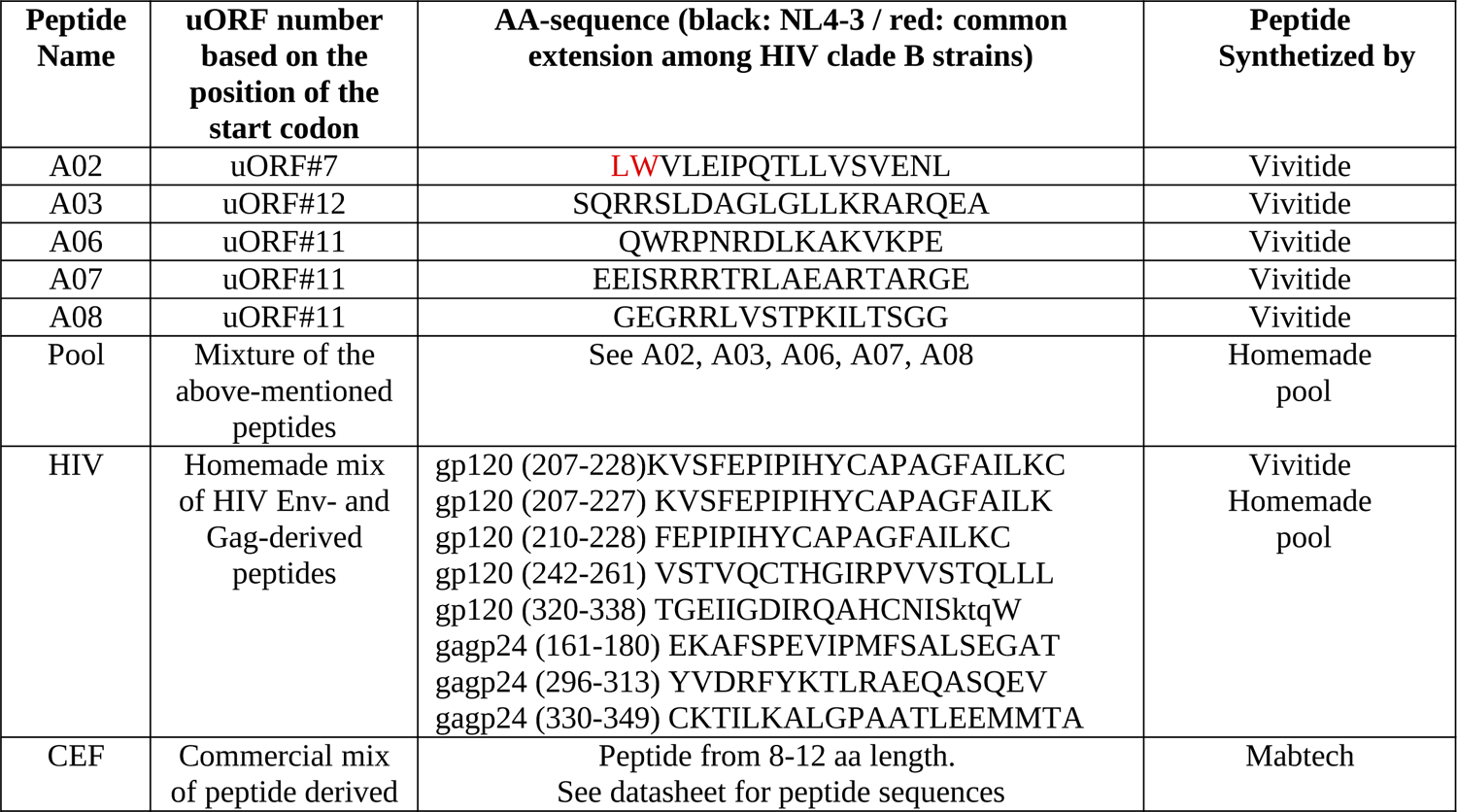
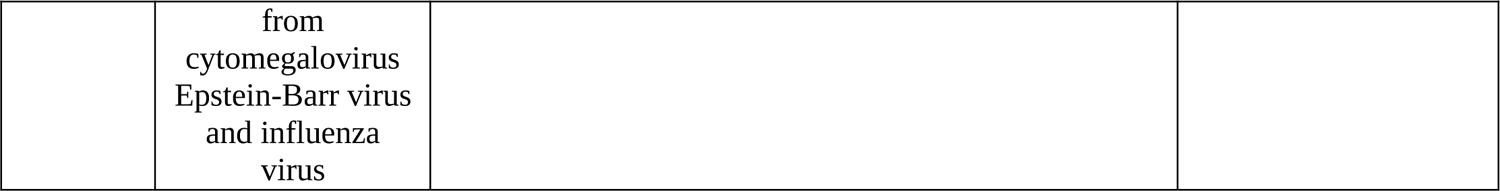

#### Enzyme-linked immunospot (IFN-γ Elispot) assay

From 10-25 x 10^5^ cells/well were seeded Elispot plates (MSIPN4550, Millipore) in IMDM supplemented with 10 % SAB, Pen/Strep, nonessential amino acids and sodium pyruvate, loaded with either 50, 10 or 2 µg/ml of uORF-derived peptides, HIV and CEF peptides, respectively and incubated at 37°C for 16h. Elispot plates were pre-coated with anti-IFNγ primary antibody and after cell-incubation IFNγ revealed using anti-IFNγ secondary antibody conjugated with biotin (both from Mabtech) as described (Cardinaud et al., 2017). The Elispot analysis was performed in technical triplicates or duplicates. Spots were counted with the AID ELISPOT Reader according to standard protocols. Responses were considered positive when IFNγ production was superior to 20 spots/10^6^ PBMCs and at least twofold higher than background (cells loaded with the DMSO containing medium used to solubilized the peptides).

#### hSHAPE analysis

Benzoyl cyanide (BzCN) was used to acylate the 2’-hydroxyl group of the unconstrained nucleotides in the RNA structure, followed by interrogation of each nucleotide using two sets of identical but differentially labeled primers: one set (AS1 and AS2) corresponds to the sequence 5’-TCGCTTTCAAGTCCCTGTTCG-3’ (complementary to HIV-1 nt 189-209) and the second set (AS3 and AS4) correspond to 5’-TTCTTTCCCCCTGGCCTTAAC-3’ (complementary to luciferase sequence nt 395-415). One primer within each set was labeled with either VIC (AS1 & AS3) or NED (AS2 & AS4). The NED-labeled primers from each set were used to prepare a ddG sequencing ladder from the untreated RNA samples. The VIC-labeled primers were used for reverse transcription of the modified or DMSO-treated control RNAs.

Briefly, 1 pmol of *in vitro* transcribed RNA was denatured at 90°C for 2 min and then cooled on ice for 2 min, followed by the addition of excess yeast tRNA (2 μg) and RNasin (5U) in 10 μl HEPES Buffer (30 mM HEPES pH 8, 300 mM KCl, 5 mM MgCl_2_). The RNA was then folded at 37°C for 20 min and modified by 3 μl of 300 mM BzCN in DMSO one min at room temperature. After adding 82 μl of water, the chemically modified RNA was extracted (Roti-Phenol/Chloroform/Isoamyl alcohol), ethanol precipitated and resuspended in 7 μl of water. Similarly, for the control (unmodified RNA sample), 3 μl of anhydrous DMSO was added instead of BzCN and treated in the same manner.

For elongation of both the modified and control samples, 1 μl of AS1 or AS3 (1 μM) were added to the resuspended RNA and incubated at 90°C for 2 min, then cooled on ice for 2 min. Two μl of 5X RT buf er was addedto each of the samples andincubated at room temperaturefor 10 min. Reverse transcriptionwas initiated by addition of 10 μl of the elongationmix (2 μl of 5X RT Buffer, 0.6 μl of 25 mM dNTP and2U of AMV RT (Life Science)) andincubationat 42°C for 20min, 50°C for 30 min and 60°C for 10 min. For the ddG sequencingladder, 2 pmol of untreatedRNA and 1 μl of theA2 or AS4 (2 μM) were usedandtreated as above exceptfor the composition of the sequencing mix (2 μl of 5X RT Buffer, 2 μl of 100μM ddGTP, 6μl of G10 (0.25 mM dGTP, 1mM dATP, 1mM dCTP, 1mM dTTP) and2U of AMV RT (Life Science)). For eachexperiment,80μl of water were addedand cDNA was extractedusing Roti Aqua-Phenol/Chloroform/Isoamylalcohol (Carl Roth). The aqueousphase of modified or unmodified samples were pooled with the aqueousphase of the ddG sequencing lad er. The samples were then ethanol precipitated and resuspendedin 10 μl of HiDi Formamide(ABI). The sampleswere thendenaturated5min at 90°C, cooled on ice for 5 min, centrifuged, and loaded onto a 96-well plate for sequencing(Applied Biosystems 3130xl genetic analyser).

The electropherogramsobtainedwere analyzed with QuShape algorithm (Karabiber et al., 2013) to extract reactivity data for each sample. The mean reactivity data from three independent experimentswere ap lied as constraintsto the RNA sequencein RNAstructure (version 6.1; (Reuter and Mathews, 2010)). The dot bracket file obtained was then usedto draw the RNA 2D structure into Structure Editor graphical tool, a moduleof the RNAstructure software.

## Acknowledgments

We gratefully acknowledge support from the PSMN (Pôle Scientifique de Modélisation Numérique) of the ENS de Lyon for all computingresources.We thankall participantsof the ANRS CODEX cohortandthe NIH AIDS Research andReference Reagent Programfor providing drugs.We thank, Eléonore Pérès, Madeleine Duc-Dodon, Louis Gazzolo, Pierre Jalinot and Vincent Mocquet for kindly sharingimmortalizedT cells infected with HTLV-1. We thankall membersof theRMI2 team and Antoine Corbin for manuscriptproofreading.We thank Andrea Cimarelli, head of the LP2L team at CIRI. We also thank the Dormeur Foundation, Vaduz, for providing AID ELISPOT Reader at CIMI and Bernard Maillere (CEA) and his team for granting access to their ELISPOT reader.

## Contributions

EPR and TO conceived the initial study with help from DD. EPR, EL and DD designed all experiments with the exception of the Elispot tests presented in Figure 6B-F and Supplementary Figure 5C-F, which were conceived, designed and performed by AM, BCR and LB with technical assistance from ON, PF and IH. EL, DD and LG performed most experiments, with technical assistance from DC. EL performed most bioinformatics analyses. OL, AS, BA provided samples from HIV-1 infected individuals and technical support for Elispot experiments. TS created the transgenic cell-lines expressing flag-tagged ribosomes and performed ribosome immunoprecipitation experiments from infected cells. JCP and VVB performed SHAPE analysis on mutated reporter RNAs. CG, CD, LG and LE performed infection experiments in primary human CD4+ T cells. EPR wrote the manuscript (with the exception of the Elispot section which was written by AM) with contributions from all authors.

---

**Supplementary Figure 1.**
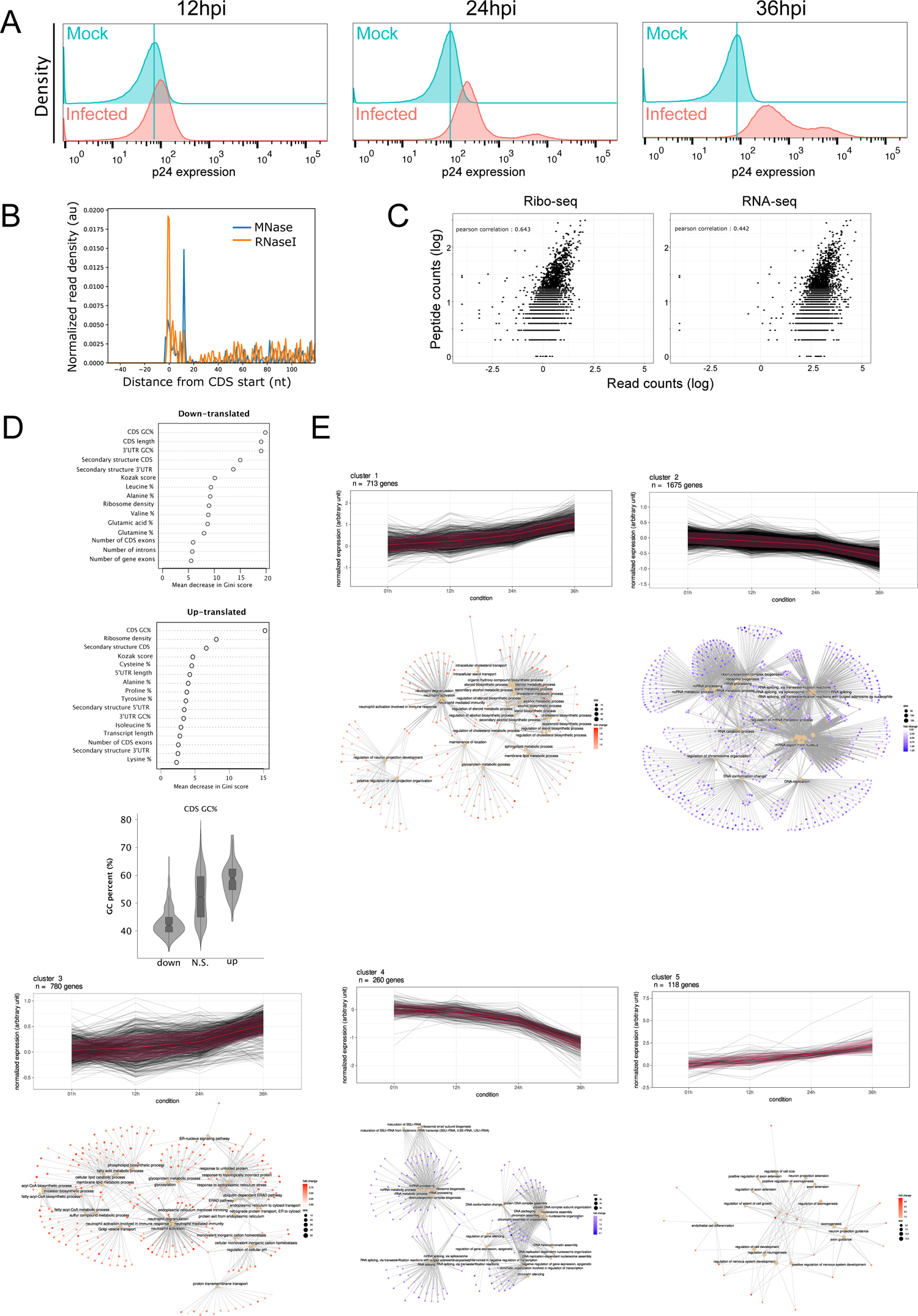
**A.** Flow cytometry analysis of p24 expression in mock (blue) or HIV-1 infected (pink) SupT1 cells at 12, 24 or 36h pi. **B.** Distribution of ribosome P-sites in the −40 to +100 nucleotide region surrounding the annotated AUG start codons of all cellular transcripts obtained from ribosome profiling libraries prepared from MNase or RNaseI-treated SupT1 cytoplasmic cell lysates. **C.** Scatter-plot of protein abundance from SupT1 cells infected with obtained by mass spectrometry (*y*-axis) from (Golumbeanu et al., 2019) against Ribo-seq (left-panel, *x*-axis) or RNA-seq (right-panel, *x*-axis) per gene read counts obtained from SupT1 cells 24 hpi from our study. **D.** Random forest analysis of features associated with down-translated (top panel), up-translated (middle panel) transcripts. (Botton panel) Violin plots of CDS GC content of down-translated (down), non-regulated (N.S.) and up-translated (up) genes at 36 hpi. **E.** Gene trajectory clustering across infection time points in SupT1 cells and its corresponding gene-ontology analysis from differentially expressed-transcripts.

**Supplementary Figure 2.**
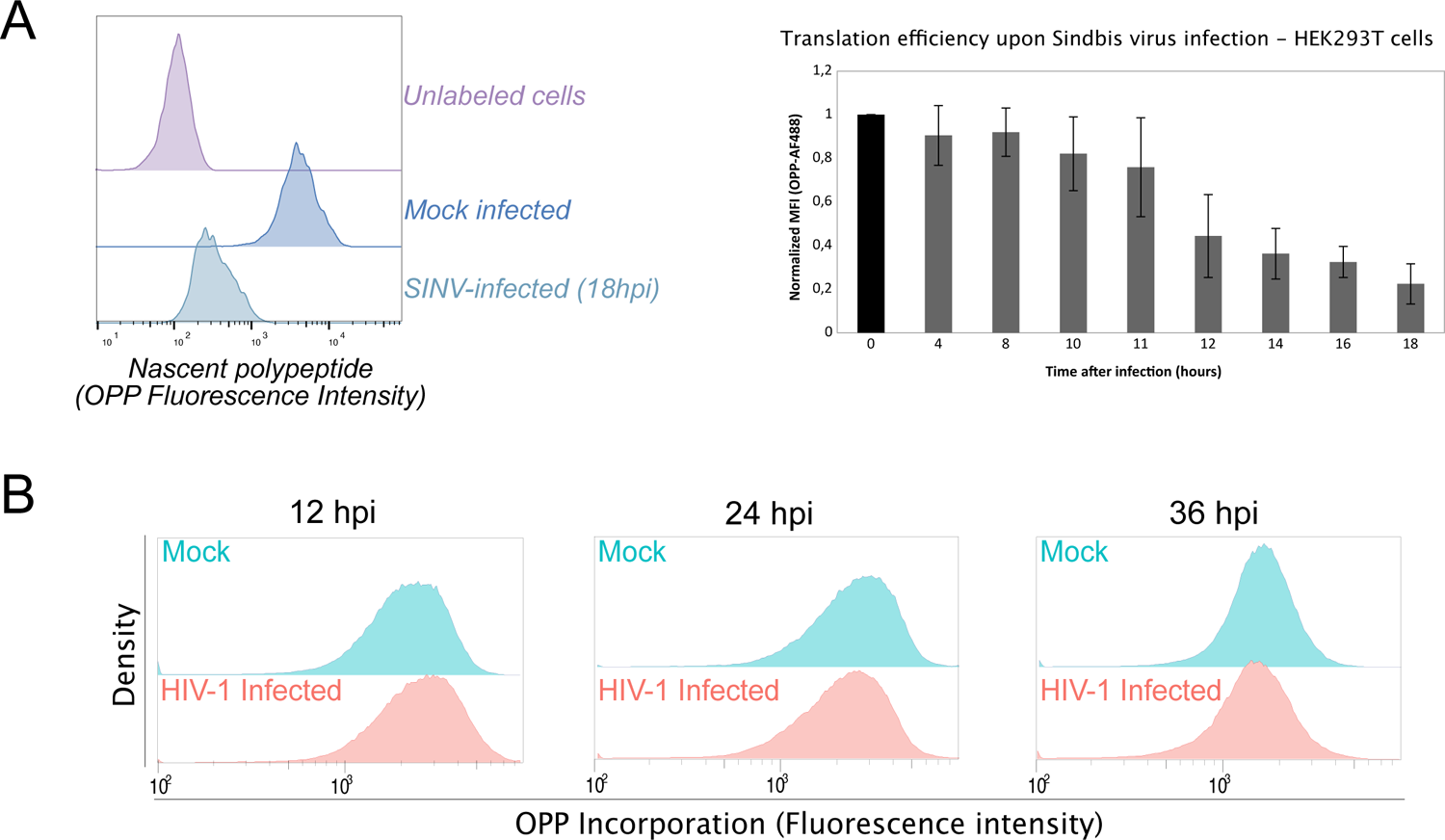
**A.** Density plot of flow cytometry signal for OPP incorporation in HEK293T cells infected with Sindbis virus (at MOI 5) from 0 to 18 hpi. **B.** Flow cytometry analysis of OPP signal in SupT1 cells mock infected or infected with HIV-1 (NL4.3 strain at MOI 5) at 12, 24 and 36 hpi (results presented correspond to gating on live cells).

**Supplementary Figure 3.**
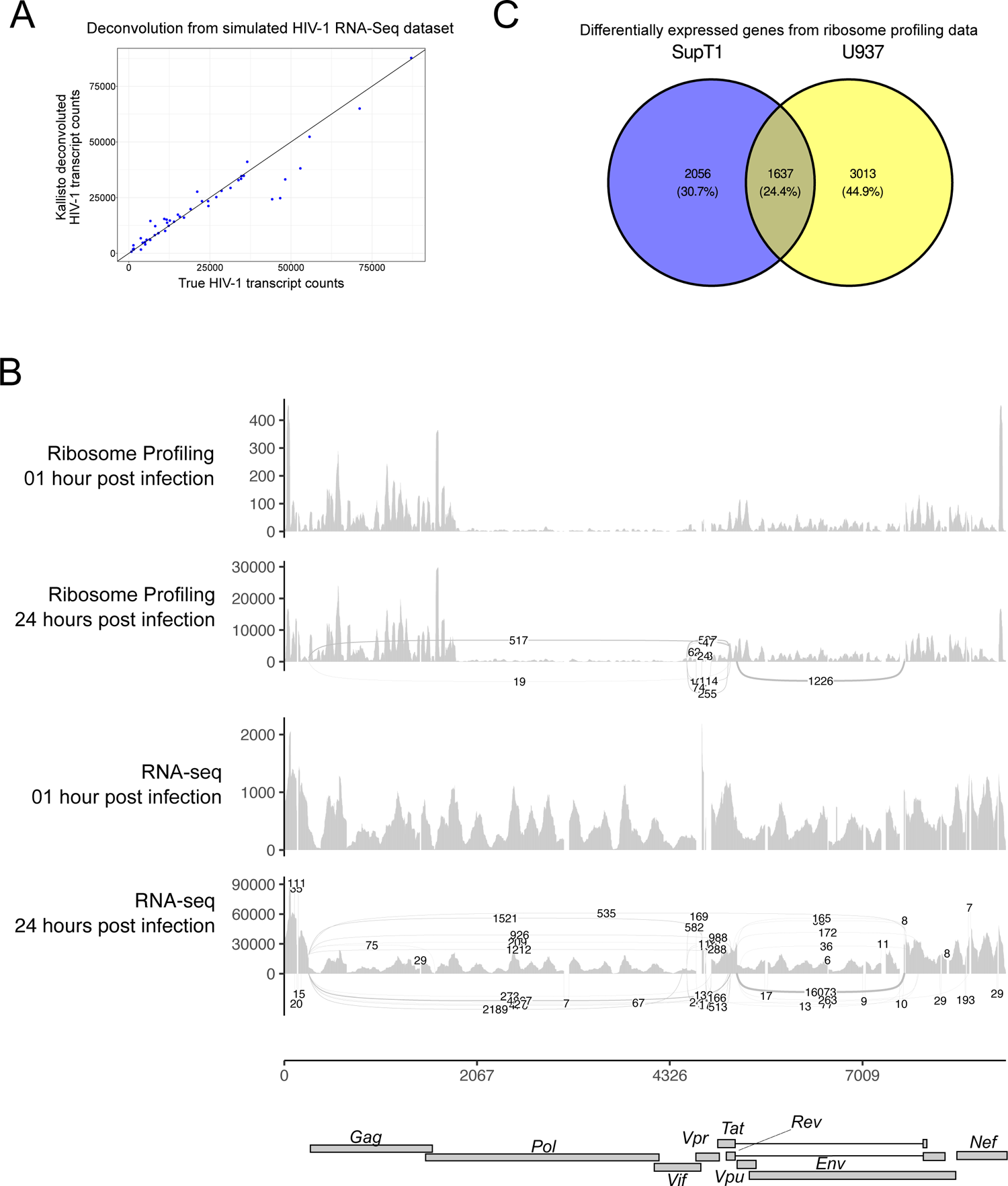
**A.** Scatter plot of a simulated HIV-1 RNA-seq dataset to test the Kallisto transcript-specific quantification pipeline based on annotated full-length transcripts obtained by Nanopore cDNA sequencing described in (Nguyen Quang et al., 2020). True HIV-1 transcript counts are presented in the *x*-axis, while Kallisto deconvoluted transcript counts are presented in the *y*-axis. **B.** Sashimi plot indicating all exon-exon junction mapping reads from our RNA-seq and Ribo-seq datasets at 1 and 24 hpi.

**Supplementary Figure 4.**
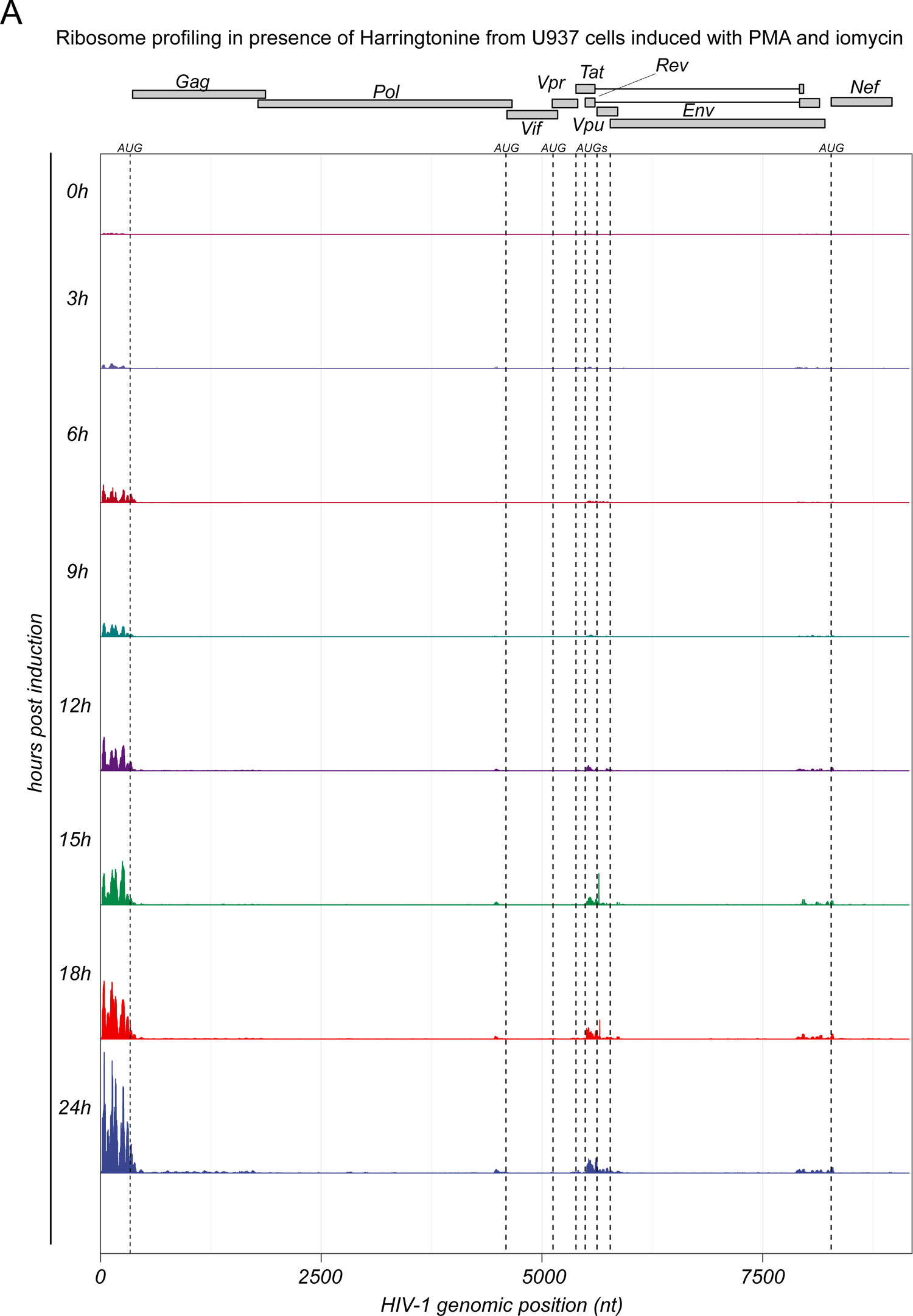
A. Distribution of ribosome profiling reads across the HIV-1 genome obtained from Harringtonine-treated U937 cells in which HIV-1 transcription was induced for 3, 6, 9, 12, 15, 18 and 24 h. The position of each viral gene and its corresponding canonical AUG start codon are depicted.

**Supplementary Figure 5.**
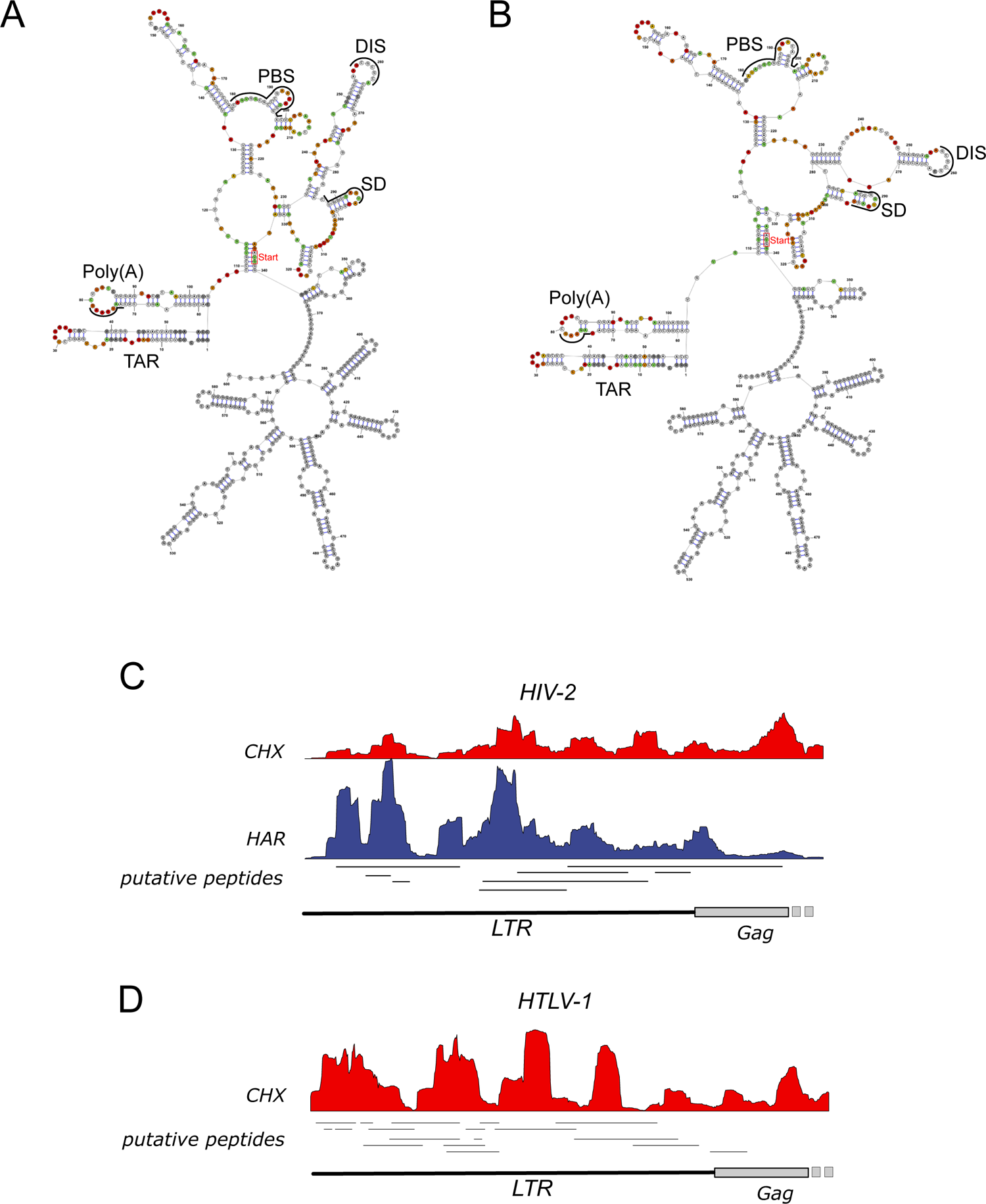
**A.** Secondary structure of the HIV-1 wild-type 5’UTR and **B.** No-uORFs 5’UTR mutant followed by the renilla luciferase coding region as obtained by SHAPE-analysis. **C.** Distribution of ribosome profiling reads across the 5’UTR of unspliced viral mRNAs in HIV-2 infected U937 cells incubated with cycloheximide or harringtonine before lysis. **D.** Distribution of ribosome profiling reads across the 5’UTR of unspliced viral mRNA of HTLV-1 in immortalized T cells obtained from HTLV-1 infected humanized mice.

**Supplementary Figure 6.**
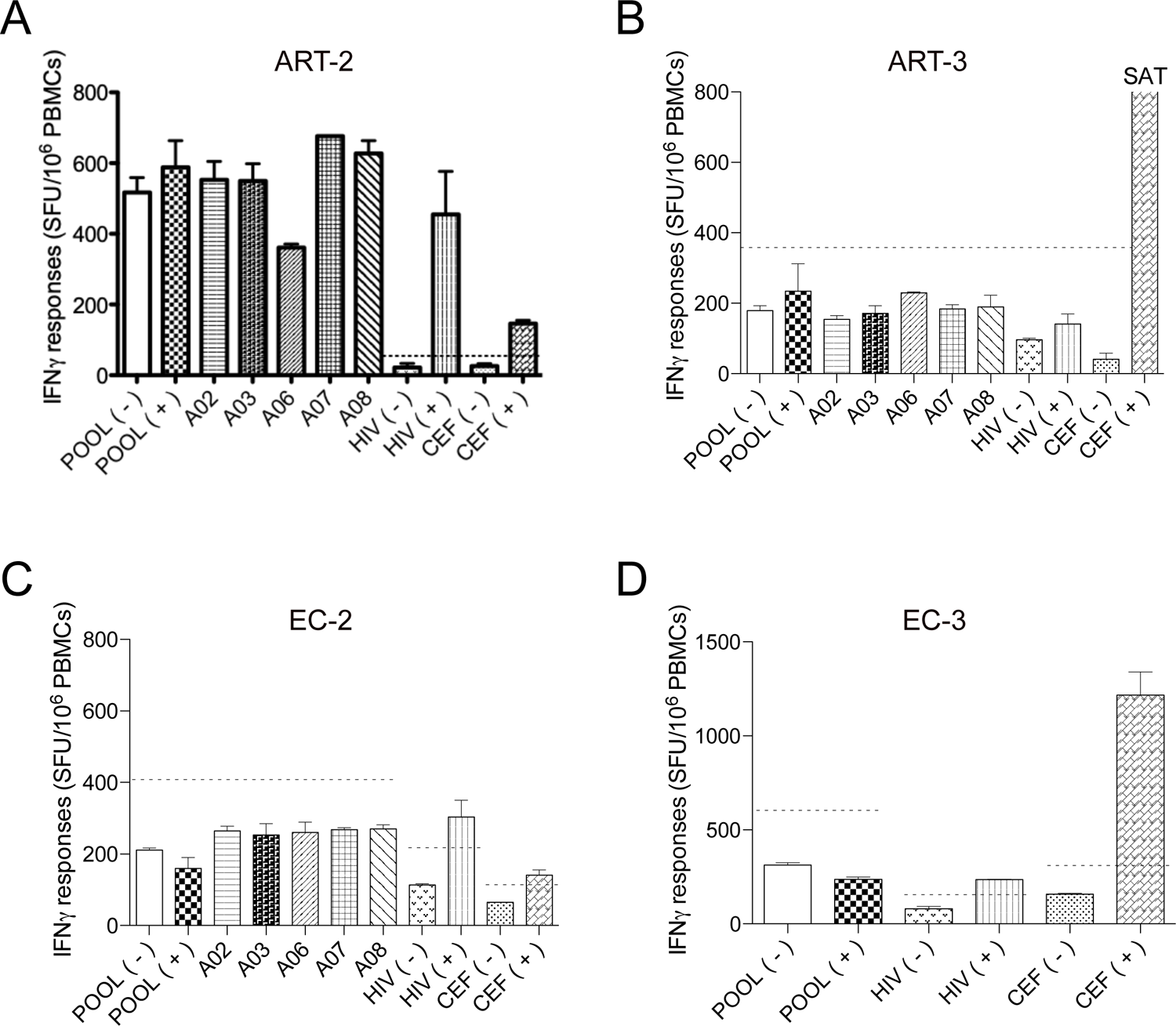
**A to D.** Four HIV infected individuals with no 5’LTR uORF-specific T cell responses. IFNγ-Elispot data from the 4 individuals, from Figure 6 **(C)**, presenting no T cell responses against 5’LTR uORF-derived peptides, expressed as spot forming units (SFU) per million PBMCs. As in Figure 6C to 6F, memory T cells from ART and EC were stimulated/expanded with the pool of uORF derived peptides (POOL), the HIV pool (HIV) or an additional control pool (CEF) composed of peptides derived from CMV, EBV and Flu viruses. **(C, D, E)**, on day 12 T cell responses against the POOL, HIV and CEF pools as well as individual 5’LTR uORF-derived peptides (A02; A03; A06; A07; A08) were assessed using IFNγ-Elispot. **(F)**, T cell responses of donor EC-3 were assessed on day 7 with the indicated pools. Since no reactivity to the pool of 5’LTR uORF-derived peptides (POOL) was detected, T cell responses to the individual peptides were not assessed on day 12 for donor EC-3. POOL (-) and POOL (+): PBMCs expanded with the pool of uORF peptides but restimulated on the day of the Elispot with medium or the POOL, respectively. A02; A03; A06; A07 and A08 name of individual uORF-derived peptides used for restimulation. HIV (-) and HIV (+): PBMCs expanded with the pool of Env- and Gag-derived peptides but restimulated for the Elispot assay with medium or the same pool of HIV peptides, respectively. CEF (-) and CEF (+) PBMCs expanded with the pool of virus-derived peptides but restimulated for the Elispot assay with medium or the same CEF pool, respectively. Responses were considered positive when IFNγ production was superior to 20 spots/10^6^ PBMCs and at least twofold higher than background production from cells restimulated with medium (dotted lines). SAT: saturated signal, where counts cannot be estimated due to overwhelm IFNγ secretion by activated T cells.

**Supplementary table 1.** Differential gene expression and translation analysis of SupT1 infected cells and U937 cells in which latent HIV-1 is induced.

**Supplementary table 2.** ORF identification from viral transcripts based on ribosome profiling reads as obtained with Ribocode.

## References

1. Adiconis, X., Borges-Rivera, D., Satija, R., DeLuca, D.S., Busby, M.A., Berlin, A.M., Sivachenko, A., Thompson, D.A., Wysoker, A., Fennell, T., et al. (2013). Comparative analysis of RNA sequencing methods for degraded or low-input samples. Nat Methods 10, 623–629. https://doi.org/10.1038/nmeth.2483.

2. Alvarez, E., Menéndez-Arias, L., and Carrasco, L. (2003). The eukaryotic translation initiation factor 4GI is cleaved by different retroviral proteases. J Virol 77, 12392–12400. https://doi.org/10.1128/jvi.77.23.12392-12400.2003.

3. Anders, S., Pyl, P.T., and Huber, W. (2015). HTSeq—a Python framework to work with high-throughput sequencing data. Bioinformatics 31, 166–169. https://doi.org/10.1093/bioinformatics/btu638.

4. Antón, L.C., and Yewdell, J.W. (2014). Translating DRiPs: MHC class I immunosurveillance of pathogens and tumors. J Leukoc Biol 95, 551–562. https://doi.org/10.1189/jlb.1113599.

5. Bansal, A., Mann, T., Sterrett, S., Peng, B.J., Bet, A., Carlson, J.M., and Goepfert, P.A. (2015). Enhanced Recognition of HIV-1 Cryptic Epitopes Restricted by HLA Class I Alleles Associated With a Favorable Clinical Outcome. JAIDS Journal of Acquired Immune Deficiency Syndromes 70, 1–8. https://doi.org/10.1097/QAI.0000000000000700.

6. Berger, C.T., Carlson, J.M., Brumme, C.J., Hartman, K.L., Brumme, Z.L., Henry, L.M., Rosato, P.C., Piechocka-Trocha, A., Brockman, M.A., Harrigan, P.R., et al. (2010). Viral adaptation to immune selection pressure by HLA class I–restricted CTL responses targeting epitopes in HIV frameshift sequences. Journal of Experimental Medicine 207, 61–75. https://doi.org/10.1084/jem.20091808.

7. Berkhout, B., Arts, K., and Abbink, T.E.M. (2011). Ribosomal scanning on the 5′-untranslated region of the human immunodeficiency virus RNA genome. Nucleic Acids Research 39, 5232– 5244. https://doi.org/10.1093/nar/gkr113.

8. Bet, A., Maze, E.A., Bansal, A., Sterrett, S., Gross, A., Graff-Dubois, S., Samri, A., Guihot, A., Katlama, C., Theodorou, I., et al. (2015). The HIV-1 Antisense Protein (ASP) induces CD8 T cell responses during chronic infection. Retrovirology 12, 15. https://doi.org/10.1186/s12977-015-0135-y.

9. Bolinger, C., Sharma, A., Singh, D., Yu, L., and Boris-Lawrie, K. (2010). RNA helicase A modulates translation of HIV-1 and infectivity of progeny virions. Nucleic Acids Res 38, 1686– 1696. https://doi.org/10.1093/nar/gkp1075.

10. Brasey, A., Lopez-Lastra, M., Ohlmann, T., Beerens, N., Berkhout, B., Darlix, J.-L., and Sonenberg, N. (2003). The leader of human immunodeficiency virus type 1 genomic RNA harbors an internal ribosome entry segment that is active during the G2/M phase of the cell cycle. J Virol 77, 3939– 3949. https://doi.org/10.1128/jvi.77.7.3939-3949.2003.

11. de Breyne, S., and Ohlmann, T. (2018). Focus on Translation Initiation of the HIV-1 mRNAs. IJMS 20, 101. https://doi.org/10.3390/ijms20010101.

12. de Breyne, S., Chamond, N., Décimo, D., Trabaud, M.-A., André, P., Sargueil, B., and Ohlmann, T. (2012). *In vitro* studies reveal that different modes of initiation on HIV-1 mRNA have different levels of requirement for eukaryotic initiation factor 4F: eIF4F requirement for HIV-1 mRNA translation. FEBS Journal 279, 3098–3111. https://doi.org/10.1111/j.1742-4658.2012.08689.x.

13. Calviello, L., Venkataramanan, S., Rogowski, K.J., Wyler, E., Wilkins, K., Tejura, M., Thai, B., Krol, J., Filipowicz, W., Landthaler, M., et al. (2021). DDX3 depletion represses translation of mRNAs with complex 5’ UTRs. Nucleic Acids Res 49, 5336–5350. https://doi.org/10.1093/nar/gkab287.

14. Cardinaud, S., Moris, A., Février, M., Rohrlich, P.-S., Weiss, L., Langlade-Demoyen, P., Lemonnier, F.A., Schwartz, O., and Habel, A. (2004). Identification of Cryptic MHC I–restricted Epitopes Encoded by HIV-1 Alternative Reading Frames. Journal of Experimental Medicine 199, 1053– 1063. https://doi.org/10.1084/jem.20031869.

15. Cardinaud, S., Consiglieri, G., Bouziat, R., Urrutia, A., Fourati, S., Malet, I., Guergnon, J., van Endert, P., Lemonnier, F.A., Appay, V., et al. (2011). CTL Escape Mediated by Proteasomal Destruction of an HIV-1 Cryptic Epitope. PLoS Pathogens 7, 15.

16. Cardinaud, S., Urrutia, A., Rouers, A., Coulon, P.-G., Kervevan, J., Richetta, C., Bet, A., Maze, E.A., Larsen, M., Iglesias, M.-C., et al. (2017). Triggering of TLR-3, −4, NOD2, and DC-SIGN reduces viral replication and increases T-cell activation capacity of HIV-infected human dendritic cells. Eur. J. Immunol. 47, 818–829. https://doi.org/10.1002/eji.201646603.

17. Casartelli, N., Guivel-Benhassine, F., Bouziat, R., Brandler, S., Schwartz, O., and Moris, A. (2010). The antiviral factor APOBEC3G improves CTL recognition of cultured HIV-infected T cells. Journal of Experimental Medicine 207, 39–49. https://doi.org/10.1084/jem.20091933.

18. Castelló, A., Franco, D., Moral-López, P., Berlanga, J.J., Alvarez, E., Wimmer, E., and Carrasco, L. (2009). HIV-1 protease inhibits Cap- and poly(A)-dependent translation upon eIF4GI and PABP cleavage. PLoS One 4, e7997. https://doi.org/10.1371/journal.pone.0007997.

19. Cenik, C., Cenik, E.S., Byeon, G.W., Grubert, F., Candille, S.I., Spacek, D., Alsallakh, B., Tilgner, H., Araya, C.L., Tang, H., et al. (2015). Integrative analysis of RNA, translation, and protein levels reveals distinct regulatory variation across humans. Genome Res 25, 1610–1621. https://doi.org/10.1101/gr.193342.115.

20. Chatel-Chaix, L., Clément, J.-F., Martel, C., Bériault, V., Gatignol, A., DesGroseillers, L., and Mouland, A.J. (2004). Identification of Staufen in the Human Immunodeficiency Virus Type 1 Gag Ribonucleoprotein Complex and a Role in Generating Infectious Viral Particles. Mol Cell Biol 24, 2637–2648. https://doi.org/10.1128/MCB.24.7.2637-2648.2004.

21. Chen, H.-H., Yu, H.-I., Yang, M.-H., and Tarn, W.-Y. (2018a). DDX3 Activates CBC-eIF3-Mediated Translation of uORF-Containing Oncogenic mRNAs to Promote Metastasis in HNSCC. Cancer Res 78, 4512–4523. https://doi.org/10.1158/0008-5472.CAN-18-0282.

22. Chen, S., Zhou, Y., Chen, Y., and Gu, J. (2018b). fastp: an ultra-fast all-in-one FASTQ preprocessor. Bioinformatics 34, i884–i890. https://doi.org/10.1093/bioinformatics/bty560.

23. Choudhary, S., Li, W., and D Smith, A. (2020). Accurate detection of short and long active ORFs using Ribo-seq data. Bioinformatics 36, 2053–2059. https://doi.org/10.1093/bioinformatics/btz878.

24. Clamer, M., Tebaldi, T., Lauria, F., Bernabò, P., Gómez-Biagi, R.F., Marchioretto, M., Kandala, D.T., Minati, L., Perenthaler, E., Gubert, D., et al. (2018). Active Ribosome Profiling with RiboLace. Cell Rep 25, 1097–1108.e5. https://doi.org/10.1016/j.celrep.2018.09.084.

25. Coulon, P.-G., Richetta, C., Rouers, A., Blanchet, F.P., Urrutia, A., Guerbois, M., Piguet, V., Theodorou, I., Bet, A., Schwartz, O., et al. (2016). HIV-Infected Dendritic Cells Present Endogenous MHC Class II–Restricted Antigens to HIV-Specific CD4+ T Cells. J Immunol 197, 517–532. https://doi.org/10.4049/jimmunol.1600286.

26. Cullen, B.R. (2003). Nuclear RNA export. J Cell Sci 116, 587–597. https://doi.org/10.1242/jcs.00268.

27. Daudé, C., Décimo, D., Trabaud, M.-A., André, P., Ohlmann, T., and de Breyne, S. (2016). HIV-1 sequences isolated from patients promote expression of shorter isoforms of the Gag polyprotein. Arch Virol 161, 3495–3507. https://doi.org/10.1007/s00705-016-3073-7.

28. Davis, A.J., Li, P., and Burrell, C.J. (1997). Kinetics of viral RNA synthesis following cell-to-cell transmission of human immunodeficiency virus type 1. J Gen Virol 78 ( *Pt 8*), 1897–1906. https://doi.org/10.1099/0022-1317-78-8-1897.

29. Deforges, J., de Breyne, S., Ameur, M., Ulryck, N., Chamond, N., Saaidi, A., Ponty, Y., Ohlmann, T., and Sargueil, B. (2017). Two ribosome recruitment sites direct multiple translation events within HIV1 Gag open reading frame. Nucleic Acids Res 45, 7382–7400. https://doi.org/10.1093/nar/gkx303.

30. Dick, R.A., Goh, S.L., Feigenson, G.W., and Vogt, V.M. (2012). HIV-1 Gag protein can sense the cholesterol and acyl chain environment in model membranes. Proc Natl Acad Sci U S A 109, 18761–18766. https://doi.org/10.1073/pnas.1209408109.

31. Dobin, A., Davis, C.A., Schlesinger, F., Drenkow, J., Zaleski, C., Jha, S., Batut, P., Chaisson, M., and Gingeras, T.R. (2013). STAR: ultrafast universal RNA-seq aligner. Bioinformatics 29, 15–21. https://doi.org/10.1093/bioinformatics/bts635.

32. Dugré-Brisson, S., Elvira, G., Boulay, K., Chatel-Chaix, L., Mouland, A.J., and Desgroseillers, L. (2004). Interaction of Staufen1 with the 5’ end of mRNA facilitates translation of these RNAs. 33, 4797–4812. https://doi.org/10.1093/nar/gki794.

33. Dunn, J.G., and Weissman, J.S. (2016). Plastid: nucleotide-resolution analysis of next-generation sequencing and genomics data. BMC Genomics 17, 958. https://doi.org/10.1186/s12864-016-3278-x.

34. Elliott, J.L., Eschbach, J.E., Koneru, P.C., Li, W., Puray-Chavez, M., Townsend, D., Lawson, D.Q., Engelman, A.N., Kvaratskhelia, M., and Kutluay, S.B. (2020). Integrase-RNA interactions underscore the critical role of integrase in HIV-1 virion morphogenesis. ELife 9, e54311. https://doi.org/10.7554/eLife.54311.

35. Emery, A., Zhou, S., Pollom, E., and Swanstrom, R. (2017). Characterizing HIV-1 Splicing by Using Next-Generation Sequencing. J Virol 91, e02515–16. https://doi.org/10.1128/JVI.02515-16.

36. Erhard, F., Dölken, L., Schilling, B., and Schlosser, A. (2020). Identification of the Cryptic HLA-I Immunopeptidome. Cancer Immunol Res 8, 1018–1026. https://doi.org/10.1158/2326-6066.CIR-19-0886.

37. Favard, C., Chojnacki, J., Merida, P., Yandrapalli, N., Mak, J., Eggeling, C., and Muriaux, D. (2019). HIV-1 Gag specifically restricts PI(4,5)P2 and cholesterol mobility in living cells creating a nanodomain platform for virus assembly. Science Advances https://doi.org/10.1126/sciadv.aaw8651.

38. Felber, B.K., Hadzopoulou-Cladaras, M., Cladaras, C., Copeland, T., and Pavlakis, G.N. (1989). rev protein of human immunodeficiency virus type 1 affects the stability and transport of the viral mRNA. Proc. Natl. Acad. Sci. U.S.A. 86, 1495–1499. https://doi.org/10.1073/pnas.86.5.1495.

39. Folks, T.M., Justement, J., Kinter, A., Dinarello, C.A., and Fauci, A.S. (1987). Cytokine-induced expression of HIV-1 in a chronically infected promonocyte cell line. Science 238, 800–802. https://doi.org/10.1126/science.3313729.

40. García-de-Gracia, F., Gaete-Argel, A., Riquelme-Barrios, S., Pereira-Montecinos, C., Rojas-Araya, B., Aguilera, P., Oyarzún-Arrau, A., Rojas-Fuentes, C., Acevedo, M.L., Chnaiderman, J., et al. (2021). CBP80/20-dependent translation initiation factor (CTIF) inhibits HIV-1 Gag synthesis by targeting the function of the viral protein Rev. RNA Biology 18, 745–758. https://doi.org/10.1080/15476286.2020.1832375.

41. Garcia-Moreno, M., Noerenberg, M., Ni, S., Järvelin, A.I., González-Almela, E., Lenz, C.E., Bach-Pages, M., Cox, V., Avolio, R., Davis, T., et al. (2019). System-wide Profiling of RNA-Binding Proteins Uncovers Key Regulators of Virus Infection. Molecular Cell 74, 196–211.e11. https://doi.org/10.1016/j.molcel.2019.01.017.

42. Gerashchenko, M.V., and Gladyshev, V.N. (2017). Ribonuclease selection for ribosome profiling. Nucleic Acids Res 45, e6–e6. https://doi.org/10.1093/nar/gkw822.

43. Gilmer, O., Mailler, E., Paillart, J.-C., Mouhand, A., Tisné, C., Mak, J., Smyth, R.P., Marquet, R., and Vivet-Boudou, V. (2022). Structural maturation of the HIV-1 RNA 5’ untranslated region by Pr55 Gag and its maturation products. RNA Biology 19, 191–205. https://doi.org/10.1080/15476286.2021.2021677.

44. Golumbeanu, M., Desfarges, S., Hernandez, C., Quadroni, M., Rato, S., Mohammadi, P., Telenti, A., Beerenwinkel, N., and Ciuffi, A. (2019). Proteo-Transcriptomic Dynamics of Cellular Response to HIV-1 Infection. Sci Rep 9. https://doi.org/10.1038/s41598-018-36135-3.

45. Goujon, C., Moncorgé, O., Bauby, H., Doyle, T., Ward, C.C., Schaller, T., Hué, S., Barclay, W.S., Schulz, R., and Malim, M.H. (2013). Human MX2 is an interferon-induced post-entry inhibitor of HIV-1 infection. Nature 502, 559–562. https://doi.org/10.1038/nature12542.

46. Guenther, U.-P., Weinberg, D.E., Zubradt, M.M., Tedeschi, F.A., Stawicki, B.N., Zagore, L.L., Brar, G.A., Licatalosi, D.D., Bartel, D.P., Weissman, J.S., et al. (2018). The helicase Ded1p controls use of near-cognate translation initiation codons in 5’ UTRs. Nature 559, 130–134. https://doi.org/10.1038/s41586-018-0258-0.

47. Guerrero, S., Batisse, J., Libre, C., Bernacchi, S., Marquet, R., and Paillart, J.-C. (2015). HIV-1 replication and the cellular eukaryotic translation apparatus. Viruses 7, 199–218. https://doi.org/10.3390/v7010199.

48. Harris, M.E., and Hope, T.J. (2000). RNA export: insights from viral models. Essays Biochem 36, 115–127. https://doi.org/10.1042/bse0360115.

49. Hartman, T.R., Qian, S., Bolinger, C., Fernandez, S., Schoenberg, D.R., and Boris-Lawrie, K. (2006). RNA helicase A is necessary for translation of selected messenger RNAs. Nature Structural & Molecular Biology 13, 509–516. https://doi.org/10.1038/nsmb1092.

50. Herbreteau, C.H., Weill, L., Décimo, D., Prévôt, D., Darlix, J.-L., Sargueil, B., and Ohlmann, T. (2005). HIV-2 genomic RNA contains a novel type of IRES located downstream of its initiation codon. Nat Struct Mol Biol 12, 1001–1007. https://doi.org/10.1038/nsmb1011.

51. Heyer, E.E., Ozadam, H., Ricci, E.P., Cenik, C., and Moore, M.J. (2015). An optimized kit-free method for making strand-specific deep sequencing libraries from RNA fragments. Nucleic Acids Research 43, e2–e2. https://doi.org/10.1093/nar/gku1235.

52. Ingolia, N.T., Lareau, L.F., and Weissman, J.S. (2011). Ribosome Profiling of Mouse Embryonic Stem Cells Reveals the Complexity and Dynamics of Mammalian Proteomes. Cell 147, 789–802. https://doi.org/10.1016/j.cell.2011.10.002.

53. Ingolia, N.T., Brar, G.A., Stern-Ginossar, N., Harris, M.S., Talhouarne, G.J.S., Jackson, S.E., Wills, M.R., and Weissman, J.S. (2014). Ribosome Profiling Reveals Pervasive Translation Outside of Annotated Protein-Coding Genes. Cell Rep 8, 1365–1379. https://doi.org/10.1016/j.celrep.2014.07.045.

54. Irigoyen, N., Dinan, A.M., Brierley, I., and Firth, A.E. (2018). Ribosome profiling of the retrovirus murine leukemia virus. Retrovirology 15, 10. https://doi.org/10.1186/s12977-018-0394-5.

55. Jaafar, Z.A., and Kieft, J.S. (2019). Viral RNA structure-based strategies to manipulate translation. Nature Reviews Microbiology 17, 110. https://doi.org/10.1038/s41579-018-0117-x.

56. Jacks, T., Powert, M.D., Masiarzt, F.R., Luciwt, P.A., Barrt, P.J., and Varmus, H.E. (1988). Characterization of ribosomal frameshifting in HIV-1 gag-pol expression. 4..

57. Jayaram, D.R., Frost, S., Argov, C., Liju, V.B., Anto, N.P., Muraleedharan, A., Ben-Ari, A., Sinay, R., Smoly, I., Novoplansky, O., et al. (2021). Unraveling the hidden role of a uORF-encoded peptide as a kinase inhibitor of PKCs. Proc. Natl. Acad. Sci. U.S.A. 118, e2018899118. https://doi.org/10.1073/pnas.2018899118.

58. Jiang, L., Schlesinger, F., Davis, C.A., Zhang, Y., Li, R., Salit, M., Gingeras, T.R., and Oliver, B. (2011). Synthetic spike-in standards for RNA-seq experiments. Genome Res 21, 1543–1551. https://doi.org/10.1101/gr.121095.111.

59. Karabiber, F., McGinnis, J.L., Favorov, O.V., and Weeks, K.M. (2013). QuShape: Rapid, accurate, and best-practices quantification of nucleic acid probing information, resolved by capillary electrophoresis. RNA 19, 63–73. https://doi.org/10.1261/rna.036327.112.

60. Kim, D., Paggi, J.M., Park, C., Bennett, C., and Salzberg, S.L. (2019). Graph-based genome alignment and genotyping with HISAT2 and HISAT-genotype. Nat Biotechnol 37, 907–915. https://doi.org/10.1038/s41587-019-0201-4.

61. Kim, S.Y., Byrn, R., Groopman, J., and Baltimore, D. (1989). Temporal aspects of DNA and RNA synthesis during human immunodeficiency virus infection: evidence for differential gene expression. J Virol 63, 3708–3713. https://doi.org/10.1128/JVI.63.9.3708-3713.1989.

62. Langmead, B., and Salzberg, S.L. (2012). Fast gapped-read alignment with Bowtie 2. Nat Methods 9, 357–359. https://doi.org/10.1038/nmeth.1923.

63. Lefebvre, G., Desfarges, S., Uyttebroeck, F., Muñoz, M., Beerenwinkel, N., Rougemont, J., Telenti, A., and Ciuffi, A. Analysis of HIV-1 Expression Level and Sense of Transcription by High-Throughput Sequencing of the Infected Cell. Jvi.Asm.Org.

64. Li, M., Kao, E., Gao, X., Sandig, H., Limmer, K., Pavon-Eternod, M., Jones, T.E., Landry, S., Pan, T., Weitzman, M.D., et al. (2012). Codon-usage-based inhibition of HIV protein synthesis by human schlafen 11. Nature 491, 125–128. https://doi.org/10.1038/nature11433.

65. Linsalata, A.E., He, F., Malik, A.M., Glineburg, M.R., Green, K.M., Natla, S., Flores, B.N., Krans, A., Archbold, H.C., Fedak, S.J., et al. (2019). DDX3X and specific initiation factors modulate FMR1 repeat-associated non-AUG-initiated translation. EMBO Reports 20, e47498. https://doi.org/10.15252/embr.201847498.

66. Liu, J., Xu, Y., Stoleru, D., and Salic, A. (2012). Imaging protein synthesis in cells and tissues with an alkyne analog of puromycin. PNAS 109, 413–418. https://doi.org/10.1073/pnas.1111561108.

67. Liu, S., Koneru, P.C., Li, W., Pathirage, C., Engelman, A.N., Kvaratskhelia, M., and Musier-Forsyth, K. (2021). HIV-1 integrase binding to genomic RNA 5′-UTR induces local structural changes in vitro and in virio. Retrovirology 18, 37. https://doi.org/10.1186/s12977-021-00582-0.

68. Locker, N., Chamond, N., and Sargueil, B. (2011). A conserved structure within the HIV gag open reading frame that controls translation initiation directly recruits the 40S subunit and eIF3. Nucleic Acids Research 39, 2367–2377. https://doi.org/10.1093/nar/gkq1118.

69. Mangeot, P.E., Risson, V., Fusil, F., Marnef, A., Laurent, E., Blin, J., Mournetas, V., Massouridès, E., Sohier, T.J.M., Corbin, A., et al. (2019). Genome editing in primary cells and in vivo using viral-derived Nanoblades loaded with Cas9-sgRNA ribonucleoproteins. Nature Communications 10. https://doi.org/10.1038/s41467-018-07845-z.

70. Mangeot, P.E., Guiguettaz, L., Sohier, T.J.M., and Ricci, E.P. (2021). Delivery of the Cas9/sgRNA Ribonucleoprotein Complex in Immortalized and Primary Cells via Virus-like Particles (“Nanoblades”). J Vis Exp https://doi.org/10.3791/62245.

71. Meijer, H.A., and Thomas, A.A.M. (2002). Control of eukaryotic protein synthesis by upstream open reading frames in the 5’-untranslated region of an mRNA. The Biochemical Journal 367, 1– 11. https://doi.org/10.1042/BJ20011706.

72. Merino, E.J., Wilkinson, K.A., Coughlan, J.L., and Weeks, K.M. (2005). RNA Structure Analysis at Single Nucleotide Resolution by Selective 2‘-Hydroxyl Acylation and Primer Extension (SHAPE). J. Am. Chem. Soc. 127, 4223–4231. https://doi.org/10.1021/ja043822v.

73. Miele, G., Mouland, A., Harrison, G.P., Cohen, E., and Lever, A.M. (1996). The human immunodeficiency virus type 1 5’ packaging signal structure affects translation but does not function as an internal ribosome entry site structure. J Virol 70, 944–951. https://doi.org/10.1128/jvi.70.2.944-951.1996.

74. Mills, E.W., Wangen, J., Green, R., and Ingolia, N.T. (2016). Dynamic Regulation of a Ribosome Rescue Pathway in Erythroid Cells and Platelets. Cell Reports 17, 1–10. https://doi.org/10.1016/j.celrep.2016.08.088.

75. Morris, D.R., and Geballe, A.P. (2000). Upstream open reading frames as regulators of mRNA translation. Molecular and Cellular Biology 20, 8635–8642.

76. Mouzakis, K.D., Lang, A.L., Vander Meulen, K.A., Easterday, P.D., and Butcher, S.E. (2013). HIV-1 frameshift efficiency is primarily determined by the stability of base pairs positioned at the mRNA entrance channel of the ribosome. Nucleic Acids Res 41, 1901–1913. https://doi.org/10.1093/nar/gks1254.

77. Namy, O., Moran, S.J., Stuart, D.I., Gilbert, R.J.C., and Brierley, I. (2006). A mechanical explanation of RNA pseudoknot function in programmed ribosomal frameshifting. 441, 4.

78. Nguyen Quang, N., Goudey, S., Ségéral, E., Mohammad, A., Lemoine, S., Blugeon, C., Versapuech, M., Paillart, J.-C., Berlioz-Torrent, C., Emiliani, S., et al. (2020). Dynamic nanopore long-read sequencing analysis of HIV-1 splicing events during the early steps of infection. Retrovirology 17, 25. https://doi.org/10.1186/s12977-020-00533-1.

79. Nieto-Garai, J.A., Arboleya, A., Otaegi, S., Chojnacki, J., Casas, J., Fabriàs, G., Contreras, F.-X., Kräusslich, H.-G., and Lorizate, M. (2021). Cholesterol in the Viral Membrane is a Molecular Switch Governing HIV-1 Env Clustering. Adv Sci (Weinh) 8, 2003468. https://doi.org/10.1002/advs.202003468.

80. Ocwieja, K.E., Sherrill-Mix, S., Mukherjee, R., Custers-Allen, R., David, P., Brown, M., Wang, S., Link, D.R., Olson, J., Travers, K., et al. (2012). Dynamic regulation of HIV-1 mRNA populations analyzed by single-molecule enrichment and long-read sequencing. Nucleic Acids Res 40, 10345– 10355. https://doi.org/10.1093/nar/gks753.

81. Ohlmann, T., Prévôt, D., Décimo, D., Roux, F., Garin, J., Morley, S.J., and Darlix, J.-L. (2002). In Vitro Cleavage of eIF4GI but not eIF4GII by HIV-1 Protease and its Effects on Translation in the Rabbit Reticulocyte Lysate System. Journal of Molecular Biology 318, 9–20. https://doi.org/10.1016/S0022-2836(02)00070-0.

82. Ono, A., Waheed, A.A., and Freed, E.O. (2007). Depletion of cellular cholesterol inhibits membrane binding and higher-order multimerization of human immunodeficiency virus type 1 Gag. Virology 360, 27–35. https://doi.org/10.1016/j.virol.2006.10.011.

83. Ouspenskaia, T., Law, T., Clauser, K.R., Klaeger, S., Sarkizova, S., Aguet, F., Li, B., Christian, E., Knisbacher, B.A., Le, P.M., et al. (2021). Unannotated proteins expand the MHC-I-restricted immunopeptidome in cancer. Nat Biotechnol 1–9. https://doi.org/10.1038/s41587-021-01021-3.

84. Pierre, P. (2009). Immunity and the regulation of protein synthesis: surprising connections. Current Opinion in Immunology 8.

85. Plank, T.-D.M., Whitehurst, J.T., and Kieft, J.S. (2013). Cell type specificity and structural determinants of IRES activity from the 5′ leaders of different HIV-1 transcripts. Nucleic Acids Res 41, 6698–6714. https://doi.org/10.1093/nar/gkt358.

86. Prévôt, D., Décimo, D., Herbreteau, C.H., Roux, F., Garin, J., Darlix, J.-L., and Ohlmann, T. (2003). Characterization of a novel RNA-binding region of eIF4GI critical for ribosomal scanning. The EMBO Journal 22, 1909–1921. https://doi.org/10.1093/emboj/cdg175.

87. Puray-Chavez, M., Lee, N., Tenneti, K., Wang, Y., Vuong, H.R., Liu, Y., Horani, A., Huang, T., Gunsten, S.P., Case, J.B., et al. (2020). The translational landscape of SARS-CoV-2 and infected cells (Microbiology).

88. Ramírez, F., Ryan, D.P., Grüning, B., Bhardwaj, V., Kilpert, F., Richter, A.S., Heyne, S., Dündar, F., and Manke, T. (2016). deepTools2: a next generation web server for deep-sequencing data analysis. Nucleic Acids Research 44, W160–W165. https://doi.org/10.1093/nar/gkw257.

89. Ramos, H., Monette, A., Niu, M., Barrera, A., López-Ulloa, B., Fuentes, Y., Guizar, P., Pino, K., DesGroseillers, L., Mouland, A.J., et al. (2021). The double-stranded RNA-binding protein, Staufen1, is an IRES-transacting factor regulating HIV-1 cap-independent translation initiation. Nucleic Acids Research gkab1188. https://doi.org/10.1093/nar/gkab1188.

90. Rao, S., Lungu, C., Crespo, R., Steijaert, T.H., Gorska, A., Palstra, R.-J., Prins, H.A.B., van Ijcken, W., Mueller, Y.M., van Kampen, J.J.A., et al. (2021). Selective cell death in HIV-1-infected cells by DDX3 inhibitors leads to depletion of the inducible reservoir. Nat Commun 12, 2475. https://doi.org/10.1038/s41467-021-22608-z.

91. Reuter, J.S., and Mathews, D.H. (2010). RNAstructure: software for RNA secondary structure prediction and analysis. BMC Bioinformatics 11, 129. https://doi.org/10.1186/1471-2105-11-129.

92. Ricci, E.P., Soto Rifo, R., Herbreteau, C.H., Decimo, D., and Ohlmann, T. (2008a). Lentiviral RNAs can use different mechanisms for translation initiation. Biochem Soc Trans 36, 690–693. https://doi.org/10.1042/BST0360690.

93. Ricci, E.P., Herbreteau, C.H., Decimo, D., Schaupp, A., Datta, S.A.K., Rein, A., Darlix, J.-L., and Ohlmann, T. (2008b). In vitro expression of the HIV-2 genomic RNA is controlled by three distinct internal ribosome entry segments that are regulated by the HIV protease and the Gag polyprotein. RNA 14, 1443–1455. https://doi.org/10.1261/rna.813608.

94. Ricci, E.P., Kucukural, A., Cenik, C., Mercier, B.C., Singh, G., Heyer, E.E., Ashar-Patel, A., Peng, L., and Moore, M.J. (2014). Staufen1 senses overall transcript secondary structure to regulate translation. Nat Struct Mol Biol 21, 26–35. https://doi.org/10.1038/nsmb.2739.

95. Ringeard, M., Marchand, V., Decroly, E., Motorin, Y., and Bennasser, Y. (2019). FTSJ3 is an RNA 2′-O-methyltransferase recruited by HIV to avoid innate immune sensing. Nature 565, 500–504. https://doi.org/10.1038/s41586-018-0841-4.

96. Rocchi, C., Louvat, C., Miele, A., Batisse, J., Guillon, C., Ballut, L., Lener, D., Negroni, M., Ruff, M., Gouet, P., et al. (2021). The HIV-1 Integrase C-Terminal domain induces TAR RNA structural changes promoting Tat binding (Biochemistry).

97. Sanjana, N.E., Shalem, O., and Zhang, F. (2014). Improved vectors and genome-wide libraries for CRISPR screening. Nat Methods 11, 783–784. https://doi.org/10.1038/nmeth.3047.

98. Schubert, U., Norbury, C.C., Yewdell, J.W., and Bennink, J.R. (2000). Rapid degradation of a large fraction of newly synthesized proteins by proteasomes. 404, 5.

99. Schuhmacher, J., Heidu, S., Balchen, T., Richardson, J.R., Schmeltz, C., Sonne, J., Schweiker, J., Rammensee, H.-G., Thor Straten, P., Røder, M.A., et al. (2020). Vaccination against RhoC induces long-lasting immune responses in patients with prostate cancer: results from a phase I/II clinical trial. J Immunother Cancer 8, e001157. https://doi.org/10.1136/jitc-2020-001157.

100. Sharma, A., Yilmaz, A., Marsh, K., Cochrane, A., and Boris-Lawrie, K. (2012). Thriving under Stress: Selective Translation of HIV-1 Structural Protein mRNA during Vpr-Mediated Impairment of eIF4E Translation Activity. PLoS Pathog 8, e1002612. https://doi.org/10.1371/journal.ppat.1002612.

101. Singh, G., Seufzer, B., Song, Z., Zucko, D., Heng, X., and Boris-Lawrie, K. (2022). HIV-1 hypermethylated guanosine cap licenses specialized translation unaffected by mTOR. Proc. Natl. Acad. Sci. U.S.A. 119, e2105153118. https://doi.org/10.1073/pnas.2105153118.

102. Soto-Rifo, R., Rubilar, P.S., Limousin, T., de Breyne, S., Décimo, D., and Ohlmann, T. (2012). DEAD-box protein DDX3 associates with eIF4F to promote translation of selected mRNAs: Translation initiation mediated by DDX3. The EMBO Journal 31, 3745–3756. https://doi.org/10.1038/emboj.2012.220.

103. Soto-Rifo, R., Rubilar, P.S., and Ohlmann, T. (2013). The DEAD-box helicase DDX3 substitutes for the cap-binding protein eIF4E to promote compartmentalized translation initiation of the HIV-1 genomic RNA. Nucleic Acids Research 41, 6286–6299. https://doi.org/10.1093/nar/gkt306.

104. Vallejos, M., Carvajal, F., Pino, K., Navarrete, C., Ferres, M., Huidobro-Toro, J.P., Sargueil, B., and López-Lastra, M. (2012). Functional and Structural Analysis of the Internal Ribosome Entry Site Present in the mRNA of Natural Variants of the HIV-1. PLoS ONE 7, e35031. https://doi.org/10.1371/journal.pone.0035031.

105. Ventoso, I., Blanco, R., Perales, C., and Carrasco, L. (2001). HIV-1 protease cleaves eukaryotic initiation factor 4G and inhibits cap-dependent translation. PNAS 98, 12966–12971. https://doi.org/10.1073/pnas.231343498.

106. Wei, J., Kishton J, R., Angel, M., Conn S, C., Dalla-Venezia, N., Marcel, V., Vincent, A., Catez, F., Ferré, S., Ayadi, L., et al. (2019). Ribosomal Proteins Regulate MHC Class I Peptide Generation for Immunosurveillance. Mol Cell 73, 1162–1173. doi: https://doi.org/10.1016/j.molcel.2018.12.020.

107. Weill, L., James, L., Ulryck, N., Chamond, N., Herbreteau, C.H., Ohlmann, T., and Sargueil, B. (2010). A new type of IRES within gag coding region recruits three initiation complexes on HIV-2 genomic RNA. Nucleic Acids Research 38, 1367–1381. https://doi.org/10.1093/nar/gkp1109.

108. Wilkinson, K.A., Merino, E.J., and Weeks, K.M. (2006). Selective 2′-hydroxyl acylation analyzed by primer extension (SHAPE): quantitative RNA structure analysis at single nucleotide resolution. Nat Protoc 1, 1610–1616. https://doi.org/10.1038/nprot.2006.249.

109. Winans, S., and Goff, S.P. (2020). Mutations altering acetylated residues in the CTD of HIV-1 integrase cause defects in proviral transcription at early times after integration of viral DNA. PLOS Pathogens 16, e1009147. https://doi.org/10.1371/journal.ppat.1009147.

110. Xiao, Z., Huang, R., Xing, X., Chen, Y., Deng, H., and Yang, X. (2018). De novo annotation and characterization of the translatome with ribosome profiling data. Nucleic Acids Res 46, e61. https://doi.org/10.1093/nar/gky179.

111. Yedavalli, V.S.R.K., Neuveut, C., Chi, Y.-H., Kleiman, L., and Jeang, K.-T. (2004). Requirement of DDX3 DEAD box RNA helicase for HIV-1 Rev-RRE export function. Cell 119, 381–392. https://doi.org/10.1016/j.cell.2004.09.029.

112. Yewdell, J.W. (2020). DRiPs get molecular. Current Opinion in Immunology 7.

113. Zhang, H., Wang, Y., and Lu, J. (2019). Function and Evolution of Upstream ORFs in Eukaryotes. Trends in Biochemical Sciences https://doi.org/10.1016/j.tibs.2019.03.002.

114. Zhang, P., He, D., Xu, Y., Hou, J., Pan, B.-F., Wang, Y., Liu, T., Davis, C.M., Ehli, E.A., Tan, L., et al. (2017). Genome-wide identification and differential analysis of translational initiation. Nat Commun 8, 1749. https://doi.org/10.1038/s41467-017-01981-8.

